# The tiny Cretaceous stem-bird *Oculudentavis* revealed as a bizarre lizard

**DOI:** 10.1101/2020.08.09.243048

**Authors:** Arnau Bolet, Edward L. Stanley, Juan D. Daza, J. Salvador Arias, Andrej Čerňanský, Marta Vidal-García, Aaron M. Bauer, Joseph J. Bevitt, Adolf Peretti, Susan E. Evans

**Affiliations:** Institut Català de Paleontologia, Universitat Autònoma de Barcelona. Barcelona, Spain; School of Earth Sciences, University of Bristol, Bristol, United Kingdom; Department of Herpetology, Florida Museum of Natural History, Gainesville, Florida, United States; Department of Biological Sciences, Sam Houston State University, Huntsville, Texas, United States; Fundación Miguel Lillo, CONICET, San Miguel de Tucumán, Argentina; Department of Ecology, Laboratory of Evolutionary Biology, Faculty of Natural Sciences, Comenius University in Bratislava, Bratislava, Slovakia.; Department of Cell Biology & Anatomy, University of Calgary, Calgary, Canada; Department of Biology and Center for Biodiversity and Ecosystem Stewardship, Villanova University, Villanova, Pennsylvania, United States.; Australian Centre for Neutron Scattering, Australian Nuclear Science and Technology Organisation, Sydney, Australia.; GRS Gemresearch Swisslab AG and Peretti Museum Foundation, Meggen, Switzerland; Department of Cell and Developmental Biology, University College London, London, United Kingdom.

## Abstract

*Oculudentavis khaungraae* was described based on a tiny skull trapped in amber. The slender tapering rostrum with retracted osseous nares, large eyes, and short vaulted braincase led to its identification as the smallest avian dinosaur on record, comparable to the smallest living hummingbirds. Despite its bird-like appearance, *Oculudentavis* showed several features inconsistent with its original phylogenetic placement. Here we describe a more complete, specimen that demonstrates *Oculudentavis* is actually a bizarre lizard of uncertain position. The new interpretation and phylogenetic placement highlights a rare case of convergent evolution rarely seen among reptiles. Our results re-affirm the importance of Myanmar amber in yielding unusual taxa from a forest ecosystem rarely represented in the fossil record.

## Introduction

In a recent paper, Xing et al. (Xing et al., 2020a) described a tiny (∼14 mm) skull (Hupoge Amber Museum, HPG-15-3) from amber deposits in northwestern Myanmar. The skull, the holotype of a new genus and species *Oculudentavis khaungraae*, has a long tapering rostrum with retracted osseous nares, a long toothed mandible with a short symphysis, a large eye supported by prominent scleral ring, an unpaired median frontal, and a triradiate postorbital. The broad, rather rounded, parietal lacks a parietal foramen and has supratemporal processes that descend vertically to meet short quadrates with well-developed lateral concavities. Xing et al.(Xing et al., 2020a) coded the characters of *Oculudentavis* into a Mesozoic avian data matrix (Atterholt et al., 2018) and found it to be a stem-avian, one node crownward of the Jurassic *Archaeopteryx*.

GRS-Ref-28627 is a second Myanmar amber specimen preserving a skull of similar length to HPG-15-3 and a partial postcranial skeleton (Figures 1, 2A–H). Like the holotype of *Oculudentavis* (Xing et al., 2020a), it has a long rostrum, a large orbit, a short postorbital region, and a long toothed mandible. Although there are some proportional differences between the skulls as preserved, the anatomical features of individual bones strongly indicate that GRS-Ref-28627 is attributable to *Oculudentavis* (Figure 2). However, many of its characters are in conflict with the interpretation of *Oculudentavis* as a stem-bird. Instead the characters indicate that *Oculudentavis* is a lizard, albeit a highly unusual one.

**Figure 1.**
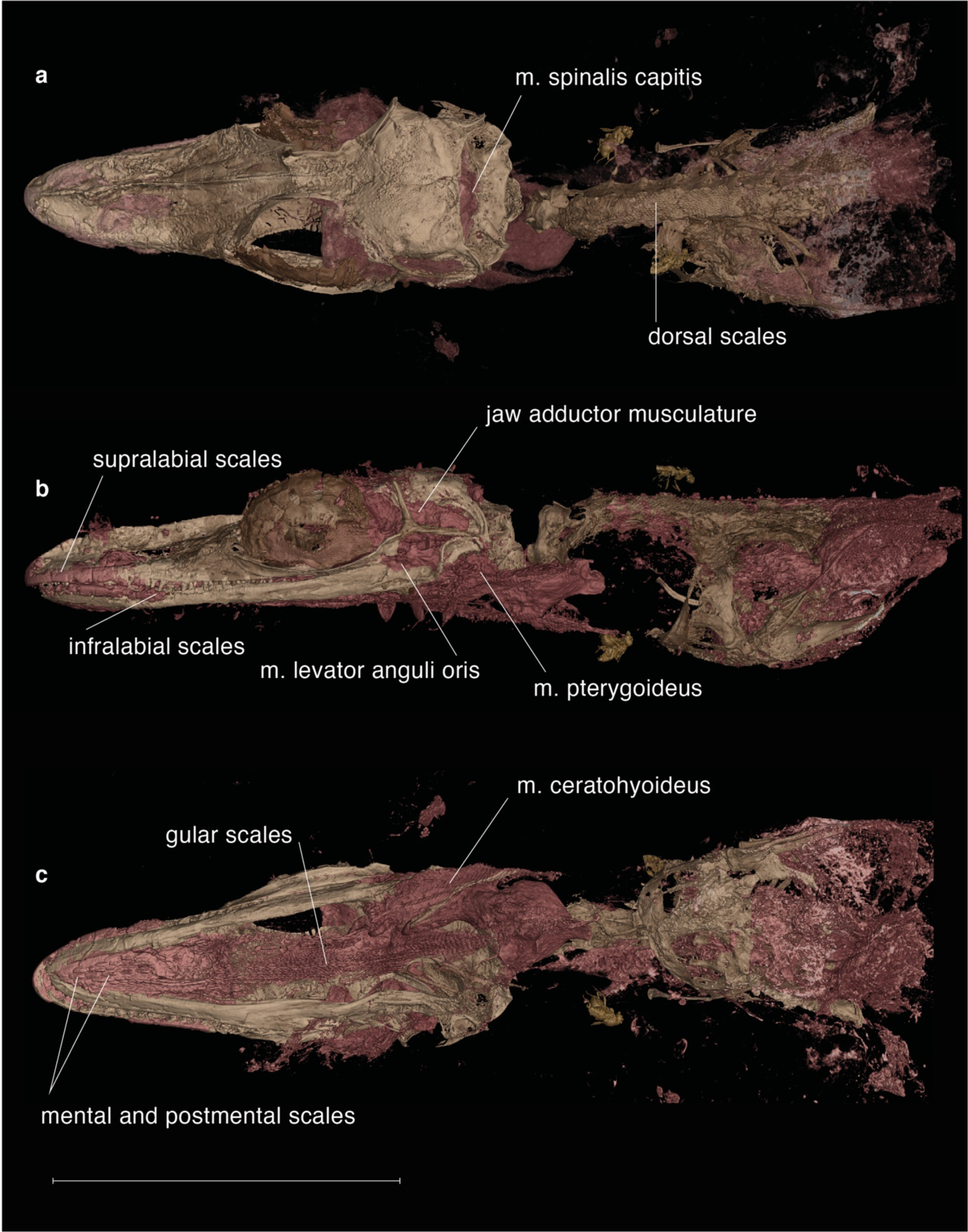
New specimen of *Ocutudentavis* sp. (GRS-Ref-286278) displaying the superb preservation of bone and soft tissue. Dorsal (a), lateral (b), and ventral (c) views. Scale bar equals 10 mm. Diptera associated were identified as Phoridae, Platypezidae, and Ceratopogonidae.

**Figure 2.**
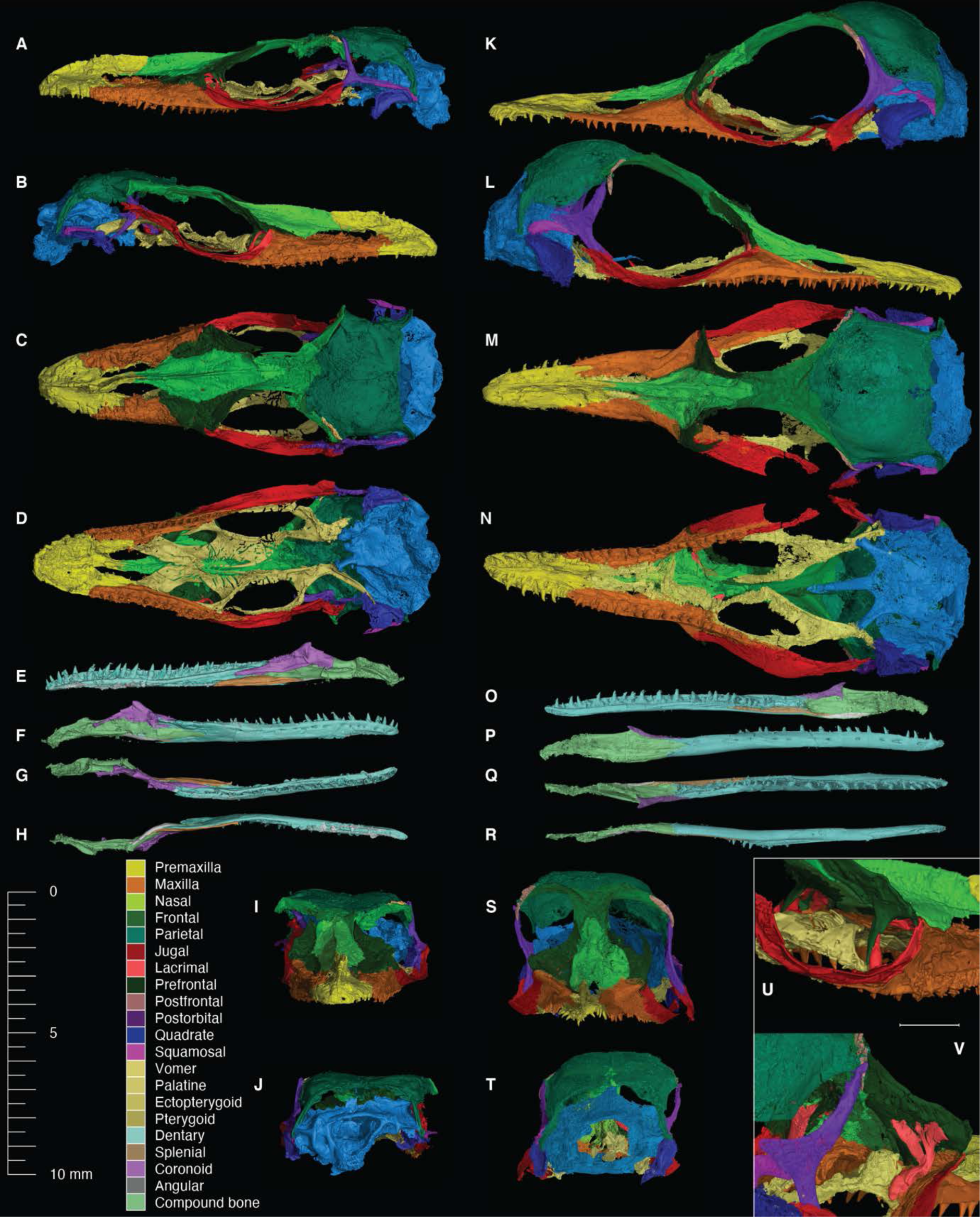
Comparison of the two specimens of *Oculudentavis*, each bone digitally segmented. Synchrotron HRCT of GRS-Ref-28627 (A–J, U) and HPG-15-3 (K–T, V). Inset shows the ring like lacrimal bone (Salmon color). Scale bar in the inset equals 1 mm.

The interpretation of HPG-15-3 as a lizard rather than a bird, has already been made (Xing et al., 2020a); one metanalysis discussed its phylogenetic placement using an ad hoc revision of diagnostic features of diapsid clades, but without testing the position of *Oculodentavis* in phylogenetic analysis (Li et al., 2020). In response, the original authors added *Oculudentavis* to a amniote data set (Pritchard and Nesbitt, 2017) and recovered *Oculodentavis* as well nested among a group of Enanthiornithes birds (O’Connor et al., 2020), arguing that placement of this taxon with squamates only occurs if all avian taxa are removed. Herein, the phylogenetic allocation of *Oculudentavis* is tested in rigorous phylogenetic analyses using data derived from the holotype of *O. khaungraae* (HPG-15-3) and additional data from the new specimen (GRS-Ref-28627). In all of our analyses (which include the amniote data set (Pritchard and Nesbitt, 2017) with some additional taxa scored by us, and a comprehensive squamate data set (Gauthier et al., 2012)), recovered *Oculodentavis a*s an squamate reptile and not as a bird. In this paper we correct the original description of this taxon, and explore the morphological data sets to determine which characteristics are convergent between *Oculodentavis* and birds.

## Results

### Systematic Paleontology

Reptilia Laurenti, 1768

Lepidosauromorpha Benton, 1983

Squamata Oppel, 1811

*Oculudentavis khaungraae* Xing et al., 2020a.

The retraction (Xing et al., 2020b) of the original description of this taxon (Xing et al., 2020a) does not affect the nomenclatural availability of *Oculudentavis khaungraae* under the International Code of Zoological Nomenclature.

**Holotype.** Hupoge Amber Museum, HPG-15-3, a complete skull preserved in amber.

**Type Locality.** Cenomanian 98.8± 0.6 Ma (Shi et al., 2012), Aung Bar mine, Tanai

Township (Myitkyina District, Hukawng Valley, Kachin province), northern Myanmar.

### Referred specimen

Peretti Museum Foundation (GRS-Ref-28627, Figures 1, 2A–H,Q, 3, Supplementary figure 1), a skull and anterior postcranial skeleton collected from amber mines in the same region as the holotype specimen. GRS-Ref-28627 is of similar size to the holotype of *Oculudentavis* (skull length ∼ 14mm). (Three-dimensional model of new specimen available at: https://tinyurl.com/vm3bayg; authors were given access to scan data of the holotype, but do not have the authority to make it publicly available).

### Emended generic diagnosis of *Oculudentavis*

Lizard characterized by the following unique combination of derived features: large unpaired median premaxilla with a long crested nasal process; long tapering rostrum composed of premaxilla, maxillae, and elongated paired nasals that slot into a triangular frontal recess; osseous nares retracted; short vaulted parietals almost or completely fused, no parietal foramen, supratemporal processes angled vertically downward; strongly triradiate postorbital with long squamosal process reaching posterior margin of parietal; vomers contact both premaxillary and maxillary shelves; large palatal fenestra between the premaxilla and the vomers; very large suborbital fenestra; ring-shaped lacrimal fully enclosing large lacrimal foramen; no palatal dentition; long slender mandible composed mainly of shallow elongate dentary; posterolingual tooth replacement; short postdentary region with coronoid bearing a low, posteriorly set process, short deep adductor fossa, and long slender retroarticular process.

### Morphological description and comparison

Like the holotype of *Oculudentavis khaungraae* (Xing et al., 2020a), the skull of GRS- Ref-28627 has a long shallow mandible of which the dentary forms the major part (∼75%), a large number of sharp, weakly pleurodont teeth (29-30 in both specimens), and a short postdentary region including a coronoid with a low, posteriorly set, coronoid eminence. The new specimen also reveals a long retroarticular process and a short, deep adductor fossa. The upper jaw comprises a unpaired median premaxilla with small teeth and a long crested nasal process (the crest was considered taphonomic in the original specimen (Xing et al., 2020a)), and a maxilla with low, medially curved facial process, a long rostral component, and a short suborbital ramus excluded from the orbital rim by the jugal. Together with paired nasals that form a rhomboid plate, these bones form a long tapering rostrum with retracted osseous nares. As in HPG-15-3, the unpaired median frontal has weak sub-olfactory processes and clasps the nasals anteriorly; the maxilla extends posteriorly only to the anterior 1/3 of the orbit; the orbit contains a large ring of “spoon-shaped” scleral ossicles that supported a large eye; and short, partially fused, parietals have a rounded lateral profile, lack a parietal foramen, and have supratemporal processes that curve ventrally rather than posteriorly to meet the short quadrates, squamosals, and short paroccipital processes of the opisthotics. The prefrontals comprise a flat anterior plate and a weakly concave orbital plate, contacting a ring-shaped lacrimal ventrally. The lacrimal completely encloses the lacrimal foramen. The postfrontal is a small, splint-like bone, lateral to the frontal and the postorbital. The postorbital is strongly triradiate with a long squamosal process that almost reaches the posterior margin of the parietal and contacts a small squamosal. There is no lower temporal bar, no quadratojugal and the postorbital process of the jugal is short. By comparison with that of HPG-15-3, the braincase of GRS-Ref-28627 is unevenly dorsoventrally compressed, so that the right side is more damaged than the left (Supplementary figure 2). Nonetheless, comparison of the two braincases shows more similarities than differences, notably the well-developed crista prootica, short crista alaris, slender basipterygoid processes, short basisphenoid, enclosed vidian canals opening posteriorly within the basisphenoid, robust parasphenoid rostrum (base only in GRS-Ref-28627), short uncrested supraoccipital with an ossified processes ascendens, and short paroccipital processes.

These similarities support the attribution of GRS-Ref-28627 to *Oculudentavis*. However, many of the features that are clearly preserved in GRS-Ref-28627 suggest it is a lizard rather than a stem bird. These include: the absence of an antorbital fenestra; the pleurodont dentition with posterolingual tooth replacement (Supplementary figure 3); a short quadrate with a lateral conch supported dorsally by a ‘hockey-stick’ shaped squamosal; a reduced quadrate-pterygoid contact; a braincase in which the metotic fissure is divided into a small ovoid lateral opening of the recessus scalae tympani and posterior vagus foramen (differentiating it from archosaurs where the metotic fissure becomes enclosed, following a totally different development (Gauthier et al., 1988)), there is an enclosed vidian canal (posterior opening within the basisphenoid), and the prootic has both an alary process and a prominent crista prootica. These features are combined with a simple primary palate in which the vomers contact both the premaxillary and maxillary shelves, and enclose a large median palatal fenestra between themselves and the premaxilla; and there is a very large suborbital fenestra. Although only part of the postcranial skeleton is preserved in GRS-Ref-28627, it shows a short neck with seven amphicoelous cervical vertebrae and atlantal arches bearing posterior zygapophyses, and a pectoral region comprising a T-shaped interclavicle, medially expanded clavicles, and a scapulocoracoid with scapular, scapulocoracoid, and primary coracoid fenestrae. In both specimens, the head (and body in GRS-Ref-28627) is covered in small scales and neither specimen preserves any trace of feathers (Figure 1, Supplementary figure 4).

**Figure 3.**
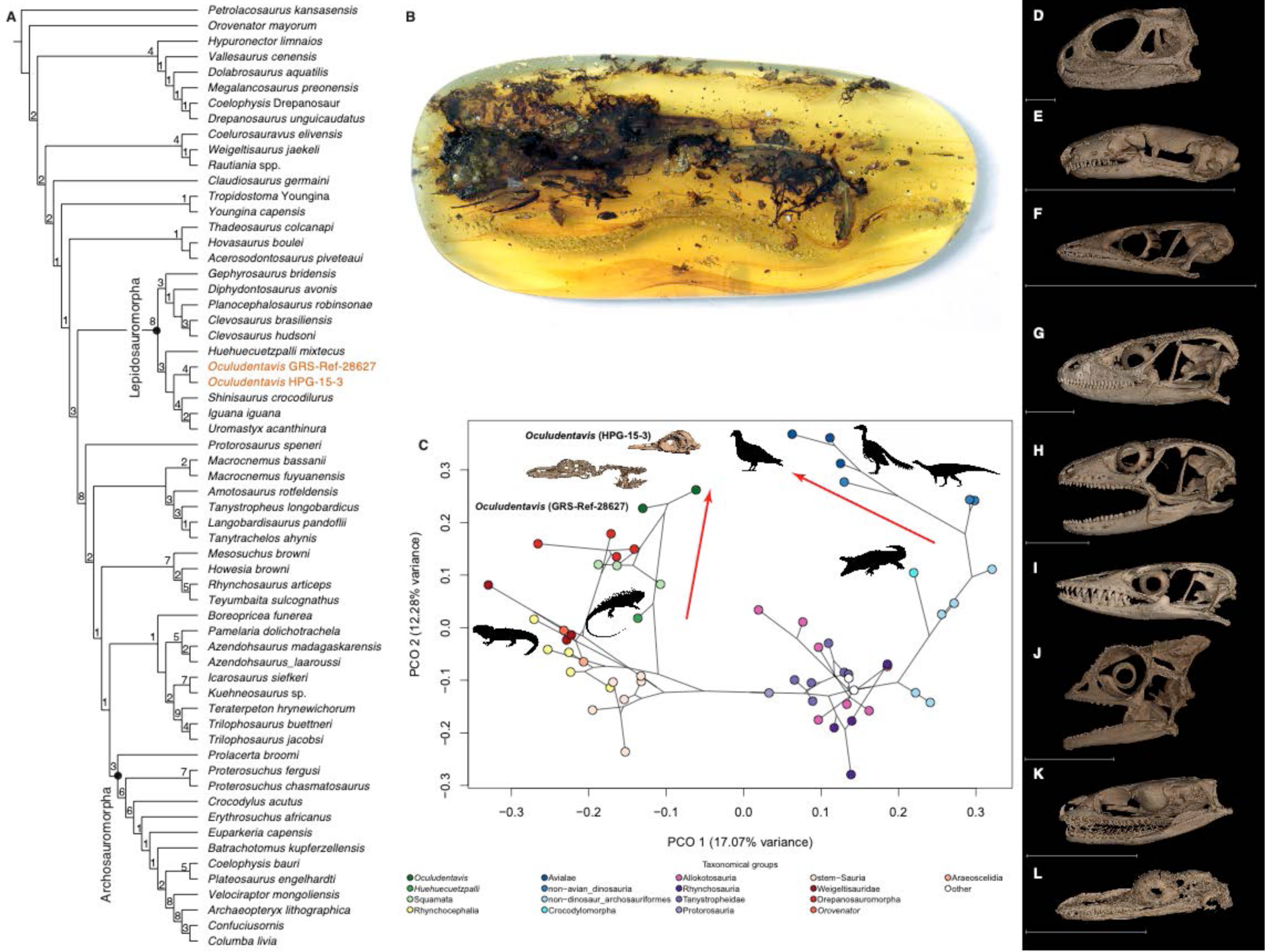
(a) Phylogenetic position of *Oculudentavis* among reptiles, values on the nodes represent Bremer support. (b) Photograph of the new specimen GRS-Ref-28627. (c) Position of *O*. *khaungraae* indicating convergence with birds according to the 2D phylomorphospace. Source of silhouettes indicated in Supplementary materials, Appendix 1. (d–l); cranial disparity of typical lepidosaurs to demonstrate the atypical skull of *O*. *khaungraae*, *Sphenodon punctatus*, Rhynchocephalia UF11978 (d); *Anelytropsis papillosus* UF-H-86708, Dibamidae (e); *Sphaerodactylus caicosensis* UF95971, Gekkota (f); *Smaug swazicus* NMB-R9201, Cordyliformes (g); *Eugongylus albofasciolatus* CAS159825, Scincidae (h); *Varanus* sp. UF71411, Varanidae (i); *Rieppeleon brevicaudatus* CAS168891, Iguania (j); *Boaedon fuliginosus* CAS85747, Serpentes (k); *Oculudentavis khaungraae* GRS-Ref-28627 (l); Scale bar equals 10 mm.

**Figure 4.**
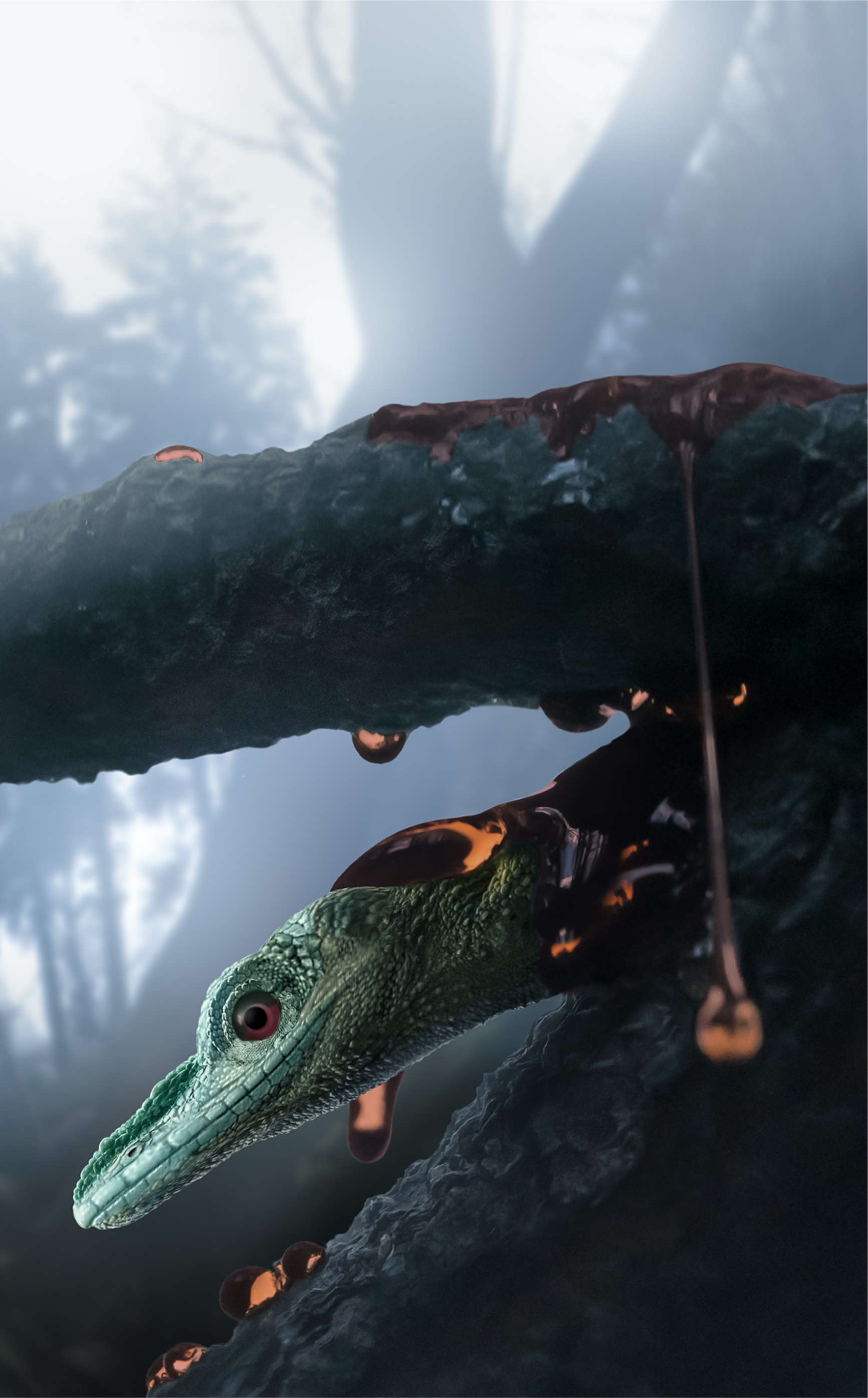
*Oculudentavis sp.* prior to being trapped in tree-resin. Scientific illustration by Stephanie Abramowicz.

Xing et al.(2020a) considered HPG-15-3 to be skeletally mature and this is probably true of the new specimen, given the tight connections between bones, the closed parietal and basicranial fontanelles, the conjoined scapula and coracoid, the sutured humeral epiphysis; and the fused dens of the axis. Any proportional differences between the specimens are therefore unlikely to be ontogenetic. HPG-15-3 appears to have a much shallower rostrum, a relatively larger eye, and a more vaulted parietal than GRS-Ref-28627, but at least some of these differences are preservational. Both specimens have undergone compression, but in different ways and in different parts of the skull. HPG-15-3 has suffered compression of the rostrum. The “dented” prefrontals and the out-turned maxillary teeth on the right side provide evidence of this, as does the fact that the teeth of the left dentary have pushed through the maxilla of that side. In an anterior view of the HPG-15-3 skull (Xing et al., 2020a), the jaw outlines are clearly distorted, and the twisting of the dentary is seen clearly in the disarticulated right mandible. In contrast, GRS-ref-28627 has experienced compression of the orbital region and posterior skull, as shown by the displacement of the tip of the left postorbital, the broken jugal, and the disparity in height between the dorsal edge of the scleral ring and the frontal. Together these points of damage exaggerate the differences in height between the rostrum, orbit, and post-orbital skull. GRS-Ref-28627 also has a taller premaxillary crest, giving the rostrum a deeper (and less bird-like) appearance. The more pronounced premaxillary crest could, potentially, be a sexually dimorphic trait. Whether the proportional differences between the skulls are fully explained by differences in size, taphonomy and sexual dimorphism, or reflect interspecific differences, is unclear. For the present, given the highly distinctive features shared by these two specimens, we refer GRS-Ref-28627 to the species *O. khaungraae*. Further specimens may clarify whether they are specifically distinct. Using modern retrodeformation methods on landmark data of the two specimens can also help to provide a proof of concept that the bird-like appearance of the HPG-15-3 skull could be an artifact of lateral compression of the snout.

GRS-Ref-28627 also preserves soft tissue (Figure 1). The head and body are covered in small, granular scales, with large and rectangular supralabial and infralabial scales, tiny scales covering the eyelid, and the nostril placed anterior to the midpoint of the retracted osseous nares (Figures 1, 4). There are no osteoderms. On the ventral surface of the head, along the midline, the epidermal scales are raised and form an evenly spaced line of short ridges. Posterior to the line of ridges, the skin is thrown into a series of narrow linear folds. This folded region underlies the hyoid ceratobranchials, and may demonstrate the resting anatomy of loose gular skin that can be inflated, for example in territorial display, in association with hyoid movements.

In its bird-like skull shape (vaulted cranium, tapering rostrum), the skull of Oculudentavis is strikingly different from any known lizard and represents a startling instance of convergent evolution. The fenestrated scapulocoracoid, hockey-stick shaped squamosal, well-developed alary process and crista prootica on the prootic, and posterolingual tooth replacement argue for a crown-group squamate position, whereas the amphicoelous vertebral centra (if not a juvenile feature) and retention of postzygapophyses on the atlas are primitive features.

### Phylogenetic and morphospace position

To test the interpretation of squamate status for *Oculudentavis*, we coded the holotype and GRS-Ref-28627 into two data sets: 1) an amniote data set that includes archosauromorphs and lepidosauromorphs (Pritchard and Nesbitt, 2017) with the addition of a non-avian theropod, a crocodylomorph, two stem-birds and a crown-bird) (Figure 3), and 2) a large morphological data matrix focused on Squamata (Gauthier et al., 2012) (Supplementary figures 5–6). The phylogenetic analysis of the amniote data set indicates unequivocally that GRS-Ref-28627 and HPG-15-3 consistently group as sister taxa, confirming the attribution of GRS-Ref-28627 to *Oculudentavis*, and that this genus clusters with squamates (Figure 3A, C). Convergence with birds can be seen in the phylomorphospace analysis using the character scores of the amniote data set, where the plot corresponding to PCO1 and PCO2 place *Oculudentavis* at the upper and central part of the lepidosauromorph morphospace, whereas birds (Figure 3C; Supplementary figures 7–8) follow the same trend for archosauromorphs. We ran a phylogenetic analysis on the squamate data set combining molecular and morphological data, and treating the data as either ordered or unordered.

A phylogenetic analysis constraining *Oculudentavis* as a bird, but without any defined location, resulted in this taxon being placed basal to a clade formed by *Archaeopteryx*+*Confusiusiornis*+*Columba*. This analysis found 12 trees that were 16 steps longer than the 48 trees of the unconstrained analysis (1159 steps [constrained] vs 1143 steps [unconstrained]). The same constrained and unconstrained analyses were run using implied weights (k=20). In these analyses the tree distortion in the the difference is 0.0140 in the case of equal weights, whereas for implied weights it is 0.0197.

## Discussion

The morphological comparison and the phylogenetic analyses strongly support the identification of GRS-Ref-28627 as a second specimen of *Oculudentavis*, and also support the interpretation of that taxon as an unusual squamate rather than a bird. However, its unusual morphology renders further attribution difficult.

Using the squamate data set, the phylogenetic placement of *Oculudentavis* within Squamata is markedly different depending on how the data is treated. If the characters are treated as ordered, the two specimens form a sister-clade to the limb-reduced, vestigial eyed, fossorial Dibamidae, near the base of Squamata (Supplementary figure 5). If characters are treated as unordered, then *Oculudentavis* is recovered within the crown-squamate clade Toxicofera (snakes, iguanians, anguimorphs (Fry et al., 2006)), on the stem of mosasaurians and snakes (Supplementary figure 6). Unfortunately, the skulls of stem-mosasaurians are poorly known (Fry et al., 2006). *Oculudentavis* shares a long ascending nasal process of the premaxilla, and retraction of the osseous nares with these taxa, but differs from them in most other respects, including the shape of the quadrate, the short parietal, absence of a parietal foramen, ventrally oriented supratemporal processes, broad vomers, absence of palatal teeth, long dentary, amphicoelous vertebrae, and fenestrated scapula (Pierce and Caldwell, 2004; Dutchak and Caldwell, 2009; Paparella et al., 2018).

It is notable that many of the characters that diagnose *Oculudentavis* are not unusual among stem birds and/or their modern descendants: large median premaxilla; long tapering rostrum composed of premaxilla, maxillae, and elongated paired nasals; osseous nares retracted; co-ossified vaulted parietals and median frontal; no parietal foramen; large orbit containing prominent scleral ossicles; primary palate without palatal teeth; short paroccipital processes; short postdentary region with coronoid bearing a low, posteriorly set process, and long slender retroarticular process. The discovery of similarity between birds and lizards, which offers tantalizing avenues for understanding the genetic drivers underpinning convergent phenotypic evolution among disparate vertebrate clades (Dutchak and Caldwell, 2009).

The convergence between *Oculudentavis* and birds is clearly depicted in the phylomorphospace plot of the amniote data set. Figure 3C shows how lepidosauromorphs and archosauromorphs separate along PCO1, whereas PCO2 highlights groupings inside these two large clades, with a tendency for more derived taxa to be situated at the top of the plot, mainly in the case of archosauromorphs. Interpreting morphospace plots is difficult for two main reasons: 1) Any given position for a taxon in morphospace (Figure 3C, Supplementary figures 7–8) is the result of a combination of phylogenetic and functional signals. However, the addition of a topology provides the necessary phylogenetic framework to confirm that the close position of *Oculudentavis* and birds in phylomorphospace is the result of morphological convergence. It is also worth noting that the lack of information on most of the postcranial skeleton of *Oculudentavis* is almost certainly exaggerating the degree of convergence. 2) Any given plot expresses a limited amount of the variance, representing the percentage corresponding to the selected axes. In our case, the 2D plot (Figure 3C), according to the first two axes, expresses about 30% of the variance, and the addition of the third axis in the 3D plot (Supplementary figure 7) raises the variance expressed to about a 39.5%. The scree data plot (Supplementary figure 9) demonstrates how the amount of variance explained by each PCO quickly decreases, the first five PCO’s accounting for roughly half of the variance, rendering the rest of PCO’s poorly informative.

Determining which characters explain the convergence in morphospace between *Oculudentavis* and birds is difficult. The observed distribution in phylomorphospace results from the sum of multiple small amounts of influence distributed across phylogenetic characters from different anatomical regions. However, a general idea of which characters are having a greater influence can be obtained using Cramér coefficients which show the strength of association between characters (or groups of characters, i.e. modules) and the PCO axes (Supplementary figure 10 shows the first 2 PCO). A detailed list and discussion of individual characters that have high or moderate Cramér coefficients is presented in *Supplementary materials*, Tables 1–4. In summary, however, the preorbital module is one of the few to show a clear signal of convergence between *Oculudentavis* and birds, notably in the concave posterior margin of the maxillary dorsal process (character 14, state 0) the presence of a lacrimal (character 24, state 0), and tooth crowns that are not attached to dentigerous bones (character 134, state 0), all primitive characters, and in the adpressed nasals separated anteriorly by the premaxilla (character 20, state 2) and absence of teeth on the bones of the palate (characters 27, 28, 30, 33-36). Note, however, that a more detailed survey of character state distribution and an increased sample might help in determining whether some of these characters underlie convergence of more inclusive groups, and if other characters might also be involved (*Supplementary materials*).

The absence of substantial postcranial material for *Oculudentavis* limits our ability to reconstruct its lifestyle. The large eyes suggest *Oculudentavis* was diurnal (Xing et al., 2020a) and, given its entrapment in amber, may have been arboreal, but this remains speculative. The mechanical advantage of a lizard jaw depends on the ratio between the distance of the jaw joint to the point of application of the adductor muscles (in-lever) and the length of the jaw as a whole (out-lever) (e.g. Stayton, 2006). Jaws in which the tooth row is of similar length to the distance between the coronoid process and the jaw articulation will deliver a stronger bite than one in which the tooth row is several times longer than the post-coronoid jaw, as it is in *Oculudentavis*. The long shallow dentary, sharp conical teeth, low coronoid process, weak mandibular symphysis, restriction of adductor muscle origin to lateral parietal margins, and the short mandibular adductor fossa are suggestive of a weak bite force. Coupled with the long retroarticular process, for the attachment of the depressor mandibulae, this implies a feeding strategy requiring fast jaw opening but limited power – perhaps for snapping at fast moving small insects (e.g. ants, or flies; Figure 1). This would be consistent with the large eyes and tapering rostrum.

With new lizard specimens emerging from the Myanmar amber each year, new specimens of *Oculudentavis* may yield additional material of the postcranial skeleton, notably the pelvic region, distal parts of the limbs, and tail, providing further eco- morphological data on this unusual lizard and further resolution of its phylogenetic position.

## Materials and Methods

### Synchrotron Scanning

Microtomographic measurements of specimen GRS-Ref-28627 were performed using the Imaging and Medical Beamline (IMBL) at the Australian Nuclear Science and Technology Organisation’s (ANSTO) Australian Synchrotron, Melbourne, Australia. For this investigation, acquisition parameters included a with a pixel size of 5.8 × 5.8 μm, monochromatic beam energy of 28 keV, a sample-to-detector distance of 100 cm and use of the “Ruby” detector consisting of a PCO.edge sCMOS camera (16-bit, 2560 × 2160 pixels) and a Nikon Makro Planar 100 mm lens coupled with a 20 μm thick Gadox/CsI(Tl)/CdWO4 scintillator screen. As the height of the specimen exceeded the detector field-of-view, the specimen was aligned axially relative to the beam and imaged using three consecutive scans, each consisting of 1800 equally-spaced angle shadow- radiographs with an exposure length of 0.50 sec, obtained every 0.10° as the sample was continuously rotated 180° about its vertical axis. Vertical translation of the specimen between tomographic scans was 11 mm. 100 dark (closed shutter) and beam profile (open shutter) images were obtained for calibration before and after shadow- radiograph acquisition. Total time for the scan was 52 min.

The raw 16-bit radiographic series were normalized relative to the beam calibration files and combined using IMBL Stitch software to yield a 32-bit series with a field-of-view of 14.8 × 29.4 mm. Reconstruction of the 3-D dataset was achieved by the filtered-back projection method and TIE-Hom algorithm phase retrieval(Paganin et al., 2002) using the CSIRO’s X-TRACT(Gureyev et al., 2011). The reconstructed volume data were rendered and visualized using VGStudio Max(Volume Graphics GmbH, 2020).

### Phylogenetic analysis

The two specimens assigned to *Oculudentavis* were retrofitted into a character matrix for lepidosaurs, including phenotypic (Gauthier et al., 2012) and molecular data (Zheng and Wiens, 2016). We were able to score 410 characters in the second specimen (GRS-Ref-28627; 67.21% scored), while only 243 in the holotype (HPG-15-3; 39.83% scored). The character scores that were available on both specimens are very similar, although part of the large amount of missing data in the holotype is due to the preservation of the specimen which makes it difficult to follow suture lines. The few character differences between the two specimens (Ch. 48, 82, 87, and 94) seem to be due to preservation. All analyses were performed with TNT (Goloboff and Catalano, 2016). We searched for the best tree over all matrices with equal weights, treating characters as either ordered (*SI Appendix*, Figures 5, 11) or ordered *SI Appendix*, Figure 6). On each run we used a script to search with new technologies (*xmult* command), starting with 20 trees of random addition, using exclusive and random sectorial searches (Goloboff, 1999), as well as 25 rounds of ratchet (Nixon, 1999) and tree drift (Goloboff, 1999), and fusing trees every five rounds. This procedure was repeated until 20 independent hits of minimal length were found. All the best trees, as well as the strict consensus were stored.

The two specimens of *Oculudentavis* were also coded into a general amniote data matrix (Pritchard and Nesbitt, 2017) to test their position in relation to both birds (represented by *Archaeopteryx*, *Confuciusornis, Columba*), and squamates. We also added the non-avian theropod *Velociraptor mongoliensis* and the crocodylomorph *Crocodylus acutus*. We were able to score 158 characters for *Oculudentavis* GRS-Ref- 28627 (51.4% of all chars), and 100 of *Oculudentavis* HPG-15-3 (32,5%). We searched for the best tree using implied weights (Goloboff, 1993; Goloboff et al., 2008; Goloboff et al., 2018), increasing the concavity value using a size of class interval of 10 from 10 to 200. As the size of the matrix is average, we used TNT (Goloboff and Catalano, 2016) with a traditional search with 200 trees of random addition (*mult* command). A majority rule tree was constructed using all the strict consensus trees of arch concavity value (Figure 4A). To test the cost of leaving *Oculudentavis* as a bird, we ran a search with equal weights in implied weights forcing *Oculudentavis* as a bird, but without any defined location and compared its length with an unconstrained tree.

### Phylomorphospace analysis

The phylomorphospace plot provides a representation of the phylogenetic relatedness of the species in our sample. It contains both the species values (the ‘tips’) and the ancestral state reconstructions for each node in the tree. All nodes and tips are connected by lines that represent the branches in the phylogeny. One advantage of phylomorphospace plots is that they make it easier to recognize whether clades and/or ecological groups plot in distinct areas of morphospace.

Phylomorphospace analyses using the morphological character matrices were run in Claddis package v. 0.2.0 (Lloyd, 2016) in R(R Core Team, 2019). One phylogeny resulting from each dataset was selected to be used in plotting phylomorphospaces. They were dated using the “equal” method of the timePaleoPhy function in the package paleotree v. 3.3.0 (Bapst, 2012). The package Plotly v. 4.9.0 (Sievert, 2018) was used to generate an interactive plot of 3D phylomorphospace. The package Geomorph v. 3.1.0 (Adams et al., 2020) was used to generate an interactive plot of 3D phylomorphospace, of which *SI Appendix*, Figure 7 is a (modified) screenshot.

The strength of association between characters and PCO axes was calculated using Cramér coefficients (See Kotrc and Knoll, 2015) as adapted in(Nordén et al., 2018). <u>Spectroscopy and stratigraphic data.</u> The Fourier-transform infrared (FTIR) spectrum of the specimen was recorded and compared to a number of reference specimens extracted directly from the Aung Bar mine by GRS staff, and to FTIR spectra of specimens obtained from other mines and localities across Myanmar. Likewise, the inclusion pattern (banding with bubble flows, etc.) was compared to other self-collected Aung Bar materials. FTIR matrix and inclusion analyses conclusively verify the indicated origin of the material from the Aung Bar mine.

GRS representatives visited the mine shaft from which the specimen was recovered, and a GoPro recording was made of the trip down the mine to the approximate location from which the specimen was found within the 1-2-meter-thick amber-containing rock layer. A miner was commissioned to recover a sample with GoPro recording of this amber-bearing rock for archiving and further analysis. Stratigraphic analysis of the entire mine column is yet to be conducted.

## Acknowledgments

We are grateful to the Peretti Museum Foundation for access to their collection of fossils in amber from Myanmar. We thank Jessica A. Maisano and Matthew Colbert from the UTCT – University of Texas – High-Resolution X-Ray CT Facility for the scanning of the referred specimen. We thank also to Lida Xing, Jingmai O’Connor for sharing the data of the Holotype, and to Gang Li at the Beijing Synchrotron Radiation Facility for facilitate the transfer of the large data set. We thank Thomas L. Stubbs for his help in calculating Cramér coefficients and for sharing the relevant R script. Special thanks to Stephanie Abramowicz for the reconstruction of *Oculudentavis*, Monica Solórzano and Enrique Peñalver for assistance identifying the flies associated with the specimen. **Competing interests:** Authors declare no competing interests. **Data and materials availability:** All data is available in the main text or the supplementary materials.

## Additional information

### Funding

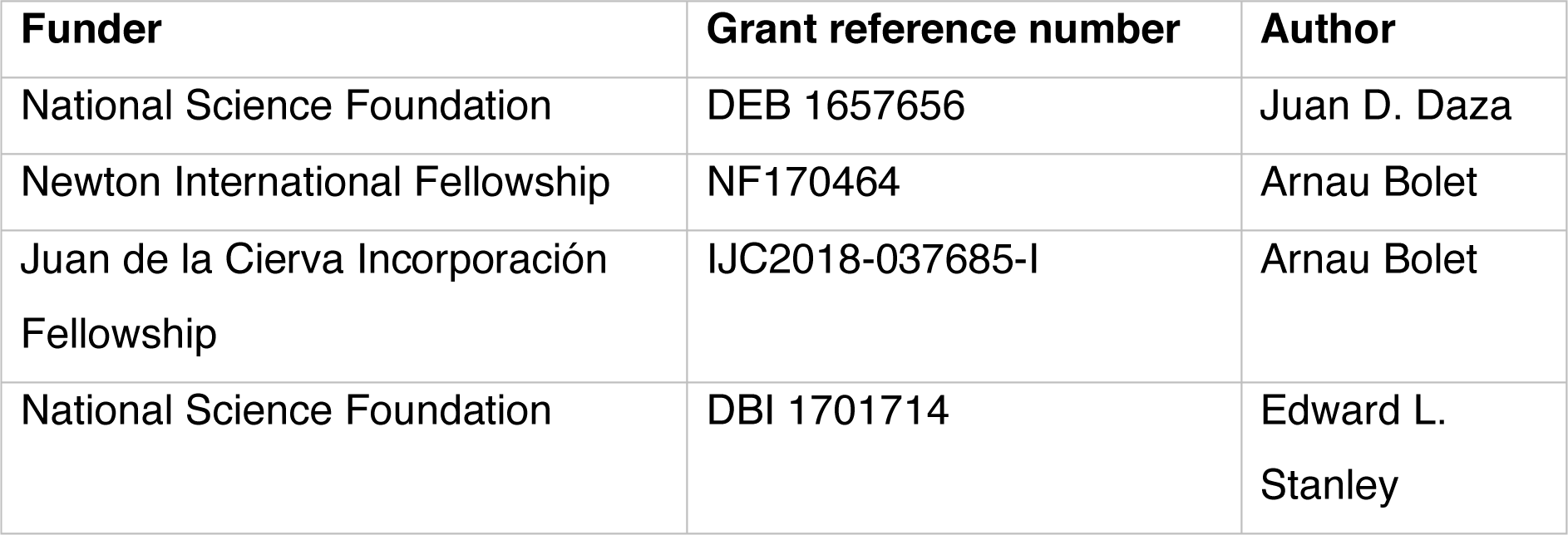

The funders had no role in study design, data collection and interpretation, or the decision to submit the work for publication.

### Author Contributions

S.E.E., A.B., E.L.S., J.D.D, A.C., A.M.B. designed research; A.B., E.L.S., J.D.D, J.S.A, A.C, MV-G, A.M.B, J.J.V performed research; A.B., E.L.S., J.D.D., J.S.A., A.C., M.V-G, A.M.B, J.J.B, A.P, S.E.E analyzed data; All authors wrote the paper; and A.P. acquired the material.

### Author ORCIDs

Arnau Bolet https://orcid.org/0000-0003-4416-4560

Edward L. Stanley https://orcid.org/0000-0001-5257-037X

Juan D. Daza https://orcid.org/0000-0002-5651-0240

J. Salvador Arias https://orcid.org/0000-0002-3717-435X

Andrej Čerňanský https://orcid.org/0000-0003-1259-2317

Marta Vidal-García https://orcid.org/0000-0001-7617-7329

Aaron M. Bauer https://orcid.org/0000-0001-6839-8025

Joseph J. Bevitt https://orcid.org/0000-0002-3502-4649

Adolf Peretti https://orcid.org/0000-0002-7080-7518

Susan E. Evans https://orcid.org/0000-0002-0799-4154

### Ethics

As researchers working on amber fossil inclusions, we follow the recommendations sent by the officers of the Society for Vertebrate Paleontology (SVP) to many Editors of scientific journals (http://vertpaleo.org/GlobalPDFS/SVP-Letter-to-Editors-FINAL.aspx).

Many of us are members of SVP, and we acknowledge their concerns as to the origin of Burmese amber material from conflict zone areas in Myanmar, human rights violations caused by its inappropriate commercialization, and the worry that some private fossil collectors are exploiting the commercial value of these fossils without considering the long-term accessibility of this material to the scientific community.

The second specimen of *Oculudentavis* (GRS-Ref-28627) and the holotype (HPG-15-3) were recovered from the same mine (Aung Bar mine, 26° 09′ N, 96° 34 E). GRS-Ref-28627 was recovered pre-conflict, in late 2017, by a local commercial miner, using local mining practices. The fossil was presented to one of the authors of this paper (AP) by local intermediaries for inspection during a humanitarian mission to the Tanai area, led by GemResearch Swiss Laboratory (GRS).

This specimen was acquired from an authorized company that exports amber pieces legally outside of Myanmar, following an ethical code that assures no violations of human rights were committed in the process of mining, and commercialization. The sample was initially loaned on consignment to GRS during a Bangkok show in 2018, for chemical, physical and spectroscopic analysis. Following these tests, the specimen was returned to Myanmar by the Myanmarese owner, and exported through a jewellery show in Yangon officially through a broker of GRS, whereupon it passed through definitive official export channels, and subsequently entered the GRS collection in Switzerland. AP strongly affirms that no funds from the sale of this amber specimen have been directed to support conflict in Kachin. The movie of the mine visit is available upon request from AP. At the time of writing, The Peretti Museum Foundation is an officially established, not-for-profit organization, founded in Switzerland by the Peretti family. The Museum promises ethical compliance in connection with the acquisition of material and supports charity projects in mining areas. The Museum actively supports research and knowledge, and provides a framework for expert collaboration and public access to all type specimens and associated content for study and publication in academic literature and other outlets. The special legal structure of the non-profit “Peretti Museum Foundation” law mandatorily guarantees under Swiss law that inventory of the foundation with GRS Reference numbers can never be lost to science.

A description and further details regarding the timing and a discussion of the military escalation in the Tanai area, and details regarding the ethical acquisition of amber pieces from Myanmar are available at the Museum website: https://www.pmf.org. The second specimen has an authenticated paper trail, including export permits from Myanmar. All documentation is available from the Peretti Museum Foundation upon request. Detail information of the ethical acquisition of amber pieces can be found in the following link: https://bit.ly/2x8gnVj.

## Additional files

### Character scores

#### Character states for the phylogenetic analysis of amniotes (Pritchard and Nesbitt, 2017) *Oculudentavis_*GRS-Ref-28627

0000??0000110111100111?0?1010000111?01??1111111?000?00?0????0?2?0??010 ??0????0110??010?01000001?000001??00?0?0??0?0?000???????????????????11110?0011011?????????????????????????????????????????????????001?0??0??0???????1????????????????001??00????0?1002?00010?10?????10?????0????????????000000??????????????0

*Oculudentavis_* HPG-15-3

000???0000110111100111?0?10?0000111?01??11111?1?000????0???????????010???????01????01??01000001?0000???????????????????????????????????????????????????????????????????????????????????????????????????????????????????????????????????????????????????????????????????????????????????????????????????????????????

*Archaeopteryx*

00010000001010100100-1-0-101000211100000???11?1-000-??00????????0??????2?????000001?110010000001000010-001???1?0???????????001-100--10020001----00110??????11001110000010-100?21100?0????1??--??11100--?001?0?0???020?1-11?200?20?10-0-1110001??0????10-00?20---10??????01--?-00????0000000-0000100??00?0100200?20?

*Confuciusornis*

00010000001010100100-1-0-101000211100???1??11?1-??????00?????????????????????000001111?1------------10-001???1?0100?00000??001-100--100???01----?0111??????10001110000010-1000211000001011??--??11100--?011?0?0??1-?0?1-1-0200?20?1?-?-1110001??00???1??1-?20---10??????01--?1??000000000----200100?00000100200?200

*Velociraptor mongoliensis*

0001100000101010010111-0-10101021110000010-11?1-000-1010????{12}?0??????????????001000110?01110000?010011-?????????????????1??001-100??11011001----0?1?1???????0????-000101101?01211000101?11----??111?0?-?001?0001?102??1?10?2?01200?0???11?00-1??00??010?00020-0010?10?--0?10?????0??000?000-0?00100??00?0?00000020?

*Crocodylus acutus*

00111000001001201111-1-0-10000021?1000001??11-1-----2010???????????????1???--001?01110?010100001010012101000010000000000111001-1000-11011001-???00110??10??10001-10001?111110120010010110?000?0-11????-?0012010??10-01(0 1)110100000000????1???100??0000011100000-??10?10?--?0110-?00?000000000-0000000-00000000200?000

*Columba livia*

00010000--1011100110-1-0-----000100000??00-1121-----0000????(1 2)?0?0??010?2???0-00?11111111------------10-00100010-1000000?01?101-1000-100---0-----001?1??1?0?10001--1200010-1010?110?0001011------111----?00110-01?1-?010-1-020-120?1-----111001--10?0-10-1-020----0010?--0---?1?00?0000000----

#### Character states for the phylogenetic analysis of lepidosaurs (Gauthier et al., 2012; Zheng and Wiens, 2016)

*Oculudentavis_*GRS-Ref-28627 110?002?011-00-0010001[0 1]100011010---100---00-0-0300200?01310--02000000020000000010102001000--0000000-0111-100?00?1313?10000010000000000000110300020102100012-010?00001???????????1000110020240????2?????????????????01011?????0-00000?10000--000120-???--02-1-01-131-0000001--001100001000000200?00????12100000?0100000110?????-00?0??0?0--?001?10?00000????100?00100010241-0?00?000100010??0?001201?010?001?1000?100000000000-030134300001110000?0000?????????????????????01002000010???????10?0???????000011?101?001010000-1?????????????????00?????????????????????????????????????????--00000000000010?00?????????????????????? *Oculudentavis_* HPG-15-3 1???0?2?0?1-00-0010001??000110??---1?0---00-0--200200?01310--02000000020000000010002000000--0100000-0111-10??00??????1???00100000000000?0110300020102100012-0100????????????????1000110020240??????????????????????010????????-???????0???--??????????????????????????0?001--??????????????????????????????????????????????????????????0--?00?????????0???????????00010241-0?00?0?0100010????????01?010??????0????00000000000-030134300001110000?0000??????????0?????????????????????????????????????????????????????????????????????????????????????????????????????????????????????????--00000000000??0?????????????????????????

### Synapomorphies for the phylogenetic analysis of lepidosaurs, ordered

For character descriptions see Gauthier et al. (2012), and apply character number + 1, as TNT uses a zero-based numbering.

Node 272 : *Oculodentavis* + DIBAMIDAE All trees:

Char. 10: 0 --> 1

Char. 27: 0 --> 1

Char. 123: 0 --> 1

Char. 193: 0 --> 2

Char. 272: 0 --> 1

Char. 311: 0 --> 1

Char. 417: 0 --> 1

Char. 426: 0 --> 1

Char. 427: 0 --> 1

Some trees: Char. 23: 0 --> 1

Char. 89: 01 --> 0

Char. 103: 0 --> 1

Char. 215: 0 --> 1

Char. 310: 0 --> 1

Char. 371: 0 --> 1

*Oculodentavis* shares with Dibamidae nine characters in all trees: Compressed premaxilla internasal process (in cross section), nasals longer than frontals, the maxilla posterior process shortens to anterior half of the orbit, fenestra ovalis opens ventrolaterally, anterior length of ectopterygoid near to or in contact with palatine, crista tuberalis reduced, maxilla tooth row length anterior to mid-orbit, marginal tooth spacing with crowns separated by large gaps, position of replacement teeth posterolingual.

In some trees, these groups share: nasals not in contact beneath the premaxillary internasal process, parietal temporal muscles originate dorsally on parietal table and supratemporal process, parietal foramen absent, vomer establish a narrow contact with the palatal shelf of the maxilla, crista interfenestralis reduced, lower dentary border of Meckel’s canal folds up to approach closely upper border to restrict canal.

Node 357 : *Oculodentavis*

All trees:

Char. 50: 0 --> 2

Char. 56: 1 --> 3

Char. 62: 0 --> 2

Char. 87: 1 --> 0

Char. 117: 0 --> 1

Char. 138: 0 --> 1

Char. 140: 0 --> 3

Char. 181: 2 --> 1

Char. 357: 0 --> 1

Char. 361: 0 --> 1

Char. 389: 2 --> 1

Char. 415: 0 --> 3

Char. 419: 3 --> 4

Char. 425: 0 --> 1

Some trees: Char. 28: 0 --> 1

Char. 35: 0 --> 1

Char. 178: 01 --> 0

Char. 327: 01 --> 0

The genus *Oculodentavis* can be diagnosed by the following 14 characters: frontal supraorbital shelf present and demarcated medially by narrow shallow longitudinal furrow often bearing a line of foramina on the dorsal surface of the frontal, frontoparietal suture deeply bowed posteriorly U or W, postfrontal with no parietal process (postfrontal subtriangular), parietals paired, maxilla narial margin rises at low angle, lacrimal foramen large, lacrimal duct enclosed entirely by the prefrontal, quadrate-pterygoid with short overlap or small lappet, dentary with reduced lateral overlap of the postdentary bones, dentary posteriormost mental foramen large, coronoid extends onto the dorsal surface of the surangular, maxillary tooth crown height reduces posteriorly, maxillary tooth count 10 or more, bases of marginal teeth expanded.

In some trees, this genus was supported by nasal terminates posterior to end of maxillary tooth row, frontals fused, lack of quadrate suprastapedial process, cultriform process long.

*Oculudentavis* GRS-Ref-28627: All trees:

Char. 86: 0 --> 1

This specimen has one autapomorphy, the postorbital extends posterior to the parietal table.

*Oculudentavis* HPG-15-3: All trees:

Char. 47: 3 --> 2

Char. 93: 0 --> 1

Some trees: Char. 81: 1 --> 0

This specimen has three autapomorphies: frontal interorbital width/frontoparietal suture width 24-26%, parietal nuchal fossa wide, and in some trees, postorbital jugal ramus extends ventral to quadrate head.

## Supplementary text

### Wildcard taxa

We identified the taxa that were more unstable in the lepidosaur data set. *Oculodentavis* was not identified as a wildcard taxon, therefore its alternative positions (as sister taxon of dibamids, ordered data) or as the sister taxon of Mosasauria+Serpentes (unordered data) can be attributed to the way that data is treated. In the analysis we did identified 13 unstable terminals (*Liushusaurus*, *Jucaraseps*, *Meyasaurus*, *Yabeinosaurus, Sakurasaurus*, *Scandensia*, *Hongshanxi*, *Hoyalacerta*,*Tchingisaurus multivagus, Gobinatus_arenosus*, *Adamisaurus_magnidentatus*, *Polyglyphanodon sternbergi*, *Gilmoreteius*). When these taxa were removed from the analysis with the characters ordered, the position of *Oculodentavis* remained stable, but the resulting tree was better resolved (*SI Appendix* Figure 10) In the same tree we also show the alternative position of the wildcard taxa:

Furthermore, the position of *Oculodentavis* close to the base of squamates is affected by the presence of dibamids. We noted that when dibamids are removed from the ordered data set, *Oculodentavis* shifts to the position recovered with the unordered data, as the sister taxon to the Mosasauria+Serpentes clade. Despite its uncertain position, we can conclude that its phylogenetic position is variable due to character codes rather than incompleteness, and that both results confirm that *Oculodentavis* is a member of Squamata and not a bird.

### Phylomorphospace analysis

Interpreting which characters or sets of characters influence each PCO axis can be challenging, mainly when using a large morphological matrix as the source data.

Cramér coefficients for PCO1 and PCO2 (Supporting Figure 6A,C), show the strength of association between characters and PCO axes. However, only a portion of the characters (about one third in the case of PCO1) provide significant values according to the calculated p-values (Supporting Figure 6B,D). Interpretation is further complicated by the fact that anatomical regions are irregular in that they contain a mixture of characters with high and low Cramér coefficients. We calculated the mean for the Cramér coefficients of all characters by anatomical region or module in an attempt to provide a comparable measure among them. According to this approach, regions that seem to have a higher number of characters strongly associated to PCO1 (Supporting Figure 6A) are the braincase and the forelimb, with particular portions of other groups of characters representing isolated high values. The order of anatomical regions according to decreasing value of the mean for PCO1 is as follows: braincase, forelimbs, skull roof and, with very similar values, the rest of regions. In PCO2 one of the few regions that clearly increase its values is the second half of characters of the preorbital region, mainly corresponding to the palate. For this axis, in order of decreasing mean value, character groups are braincase, antorbital region and hind limbs (with very similar values), and then the vertebral column, the forelimbs and skull roof and, finally, the lower jaw (note that the latter includes all characters of dentition except those for the palate).

As briefly explained in the main text, identifying the specific characters that would explain the convergence between *Oculudentavis* and birds is not an easy task. The large amount of missing data for *Oculudentavis* and, to a lesser degree, the fossil birds, results in a lack of actual support of many of these characters to the interpretation of convergence between *Oculudentavis* and birds, despite being important in the overall distribution of the rest of taxa in morphospace. Interpretation is further complicated by the fact that anatomical regions are irregular in that they contain a mixture of characters with high and low Cramér coefficients. In order to be informative to our discussion of convergence, characters need to fulfil specific criteria in relation to the distribution of their states across the morphological matrix. This requires characters that: 1) have high or moderate Cramér values; 2) are significant (p-values < 0.05) 3) are scored with the same state for at least one of the *Oculudentavis* specimens and at least one of the bird taxa and 4) are scored differently from other squamates (otherwise we would be looking at characters that show convergence between squamates and birds, not just *Oculudentavis* and birds); and 5) are not too widely distributed across the matrix.

For PCO1, although finding characters fulfilling the first three criteria is relatively easy (e.g. absence of a flattened palatal process of the premaxilla (character 7, state 0), among many others), fulfilling the fourth one is much more complicated. As explained in the main text, this is only partially accomplished by, for example, character 14, state (0), a concave posterior margin of the dorsal process of the maxilla or the presence of a lacrimal (character 24, state 0), which are shared between *Oculudentavis*, birds and some squamates. There are other characters states (e.g. absence of a cleithrum, character 208, state 1, that are of no interest to the discussion here because they are too widely distributed in the matrix. On the other hand, it is also possible that support for convergence between *Oculudentavi*s and birds is hidden among characters with an even lower Cramér value (<0.5), because we do not know at which value influence is low enough to be ignored. A more exhaustive investigation of this issue (including assessing the importance of the palate characters for PCO2 as exposed in the main text) should rely in a more exhaustive sampling of squamates and birds, and possibly in the addition of new characters more focused on tracking the evolutionary paths of lepidosaurs and archosaurs.

## Supplementary figures

**Supplementary figure 1.**
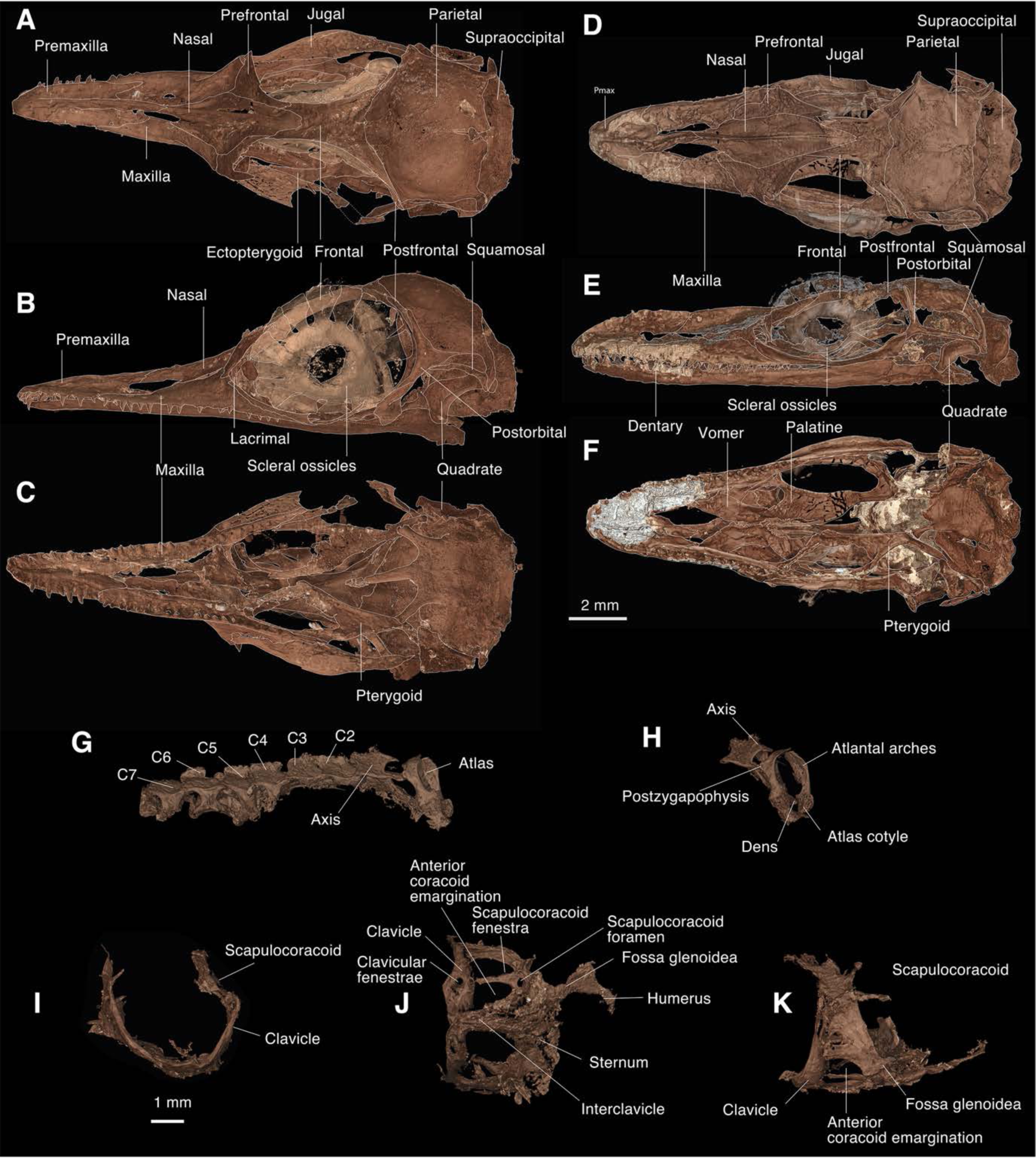
Articulated skulls of *Oculodentavis* with bone interpretation based on the digital segmentation. A–C, HPG-15-3, D–K, GRS-Ref-28627. 2 mm scale bar for skulls and 1 mm scale bar for postcranial elements. G-H, cervical vertebrae, 1- K, pectoral girdle.

**Supplementary figure 2.**
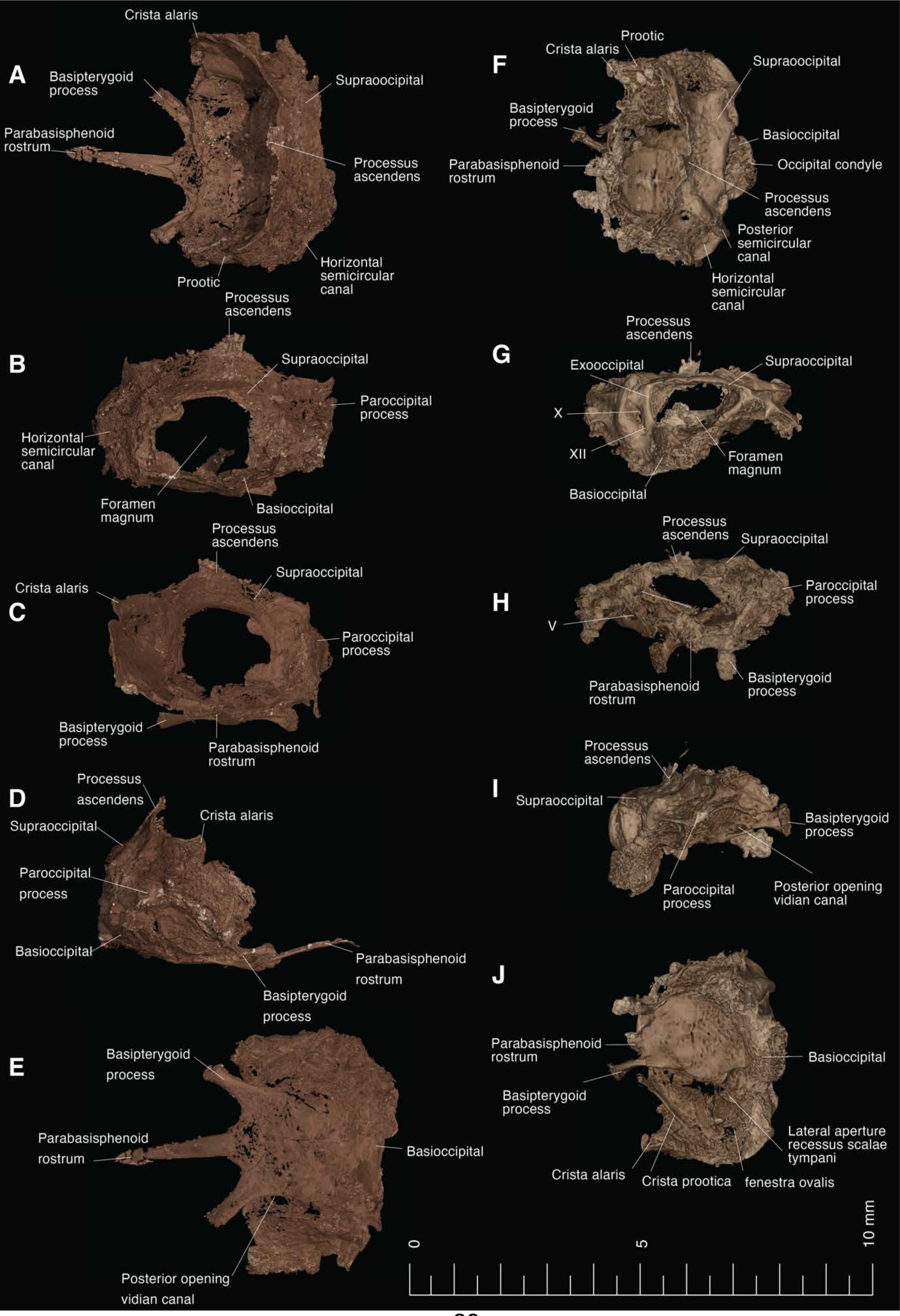
Segmented braincase of *Oculodentavis,* A-E, HPG-15-3 and F-J, GRS-Ref-28627 in dorsal, posterior, anterior, lateral (right side) and ventral views. Scale bar equals 10 mm.

**Supplementary figure 3.**
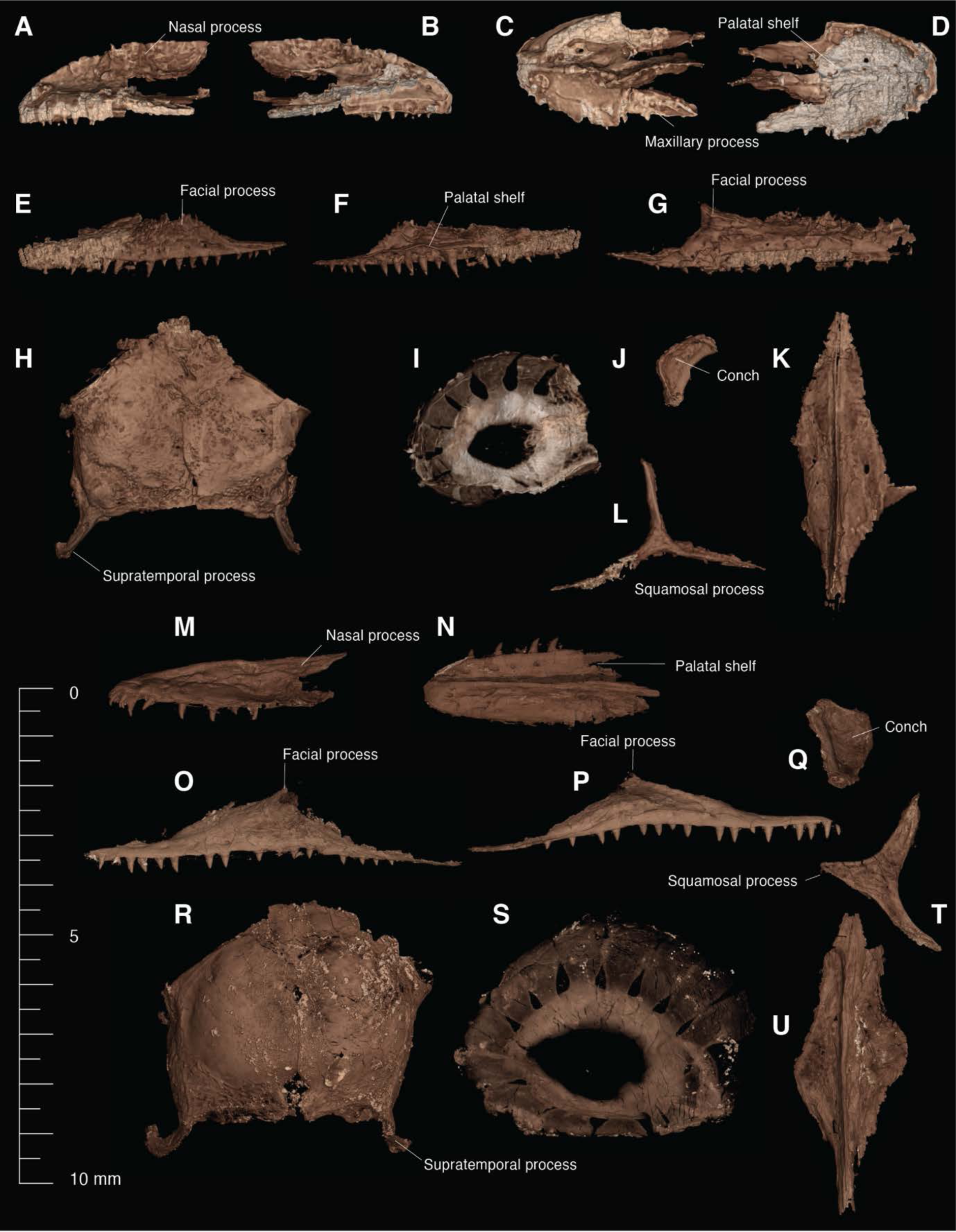
Segmented cranial bones of *Oculodentavis,* A-L, bones from GRS-Ref-28627; L-8 bones from HPG-15-3. Premaxilla A-D, L, M; maxilla E-G, N-D; parietals H, P; scleral ossicles I, Q; quadrate J, R; postorbital K, S.

**Supplementary figure 4.**
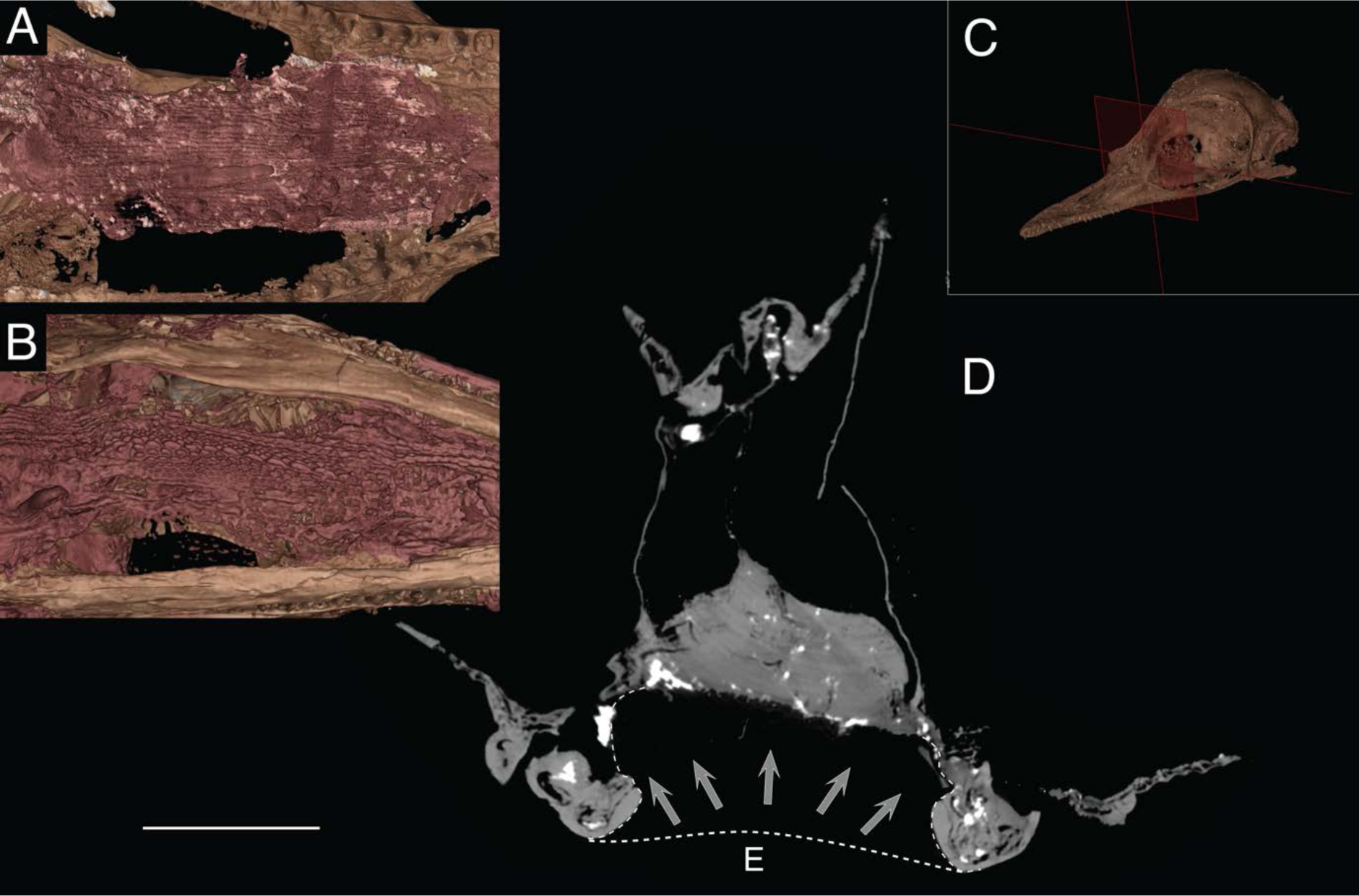
Gular scales in *Oculodentavis*, A, HPG-15-3 and B, GRS- Ref-28627. C. sagittal plane across HPG-15-3, showing the location of the tomogram (E). Dashed line shows the projected position of the scales before the skin shifted to a more dorsal position.

**Supplementary figure 5.**
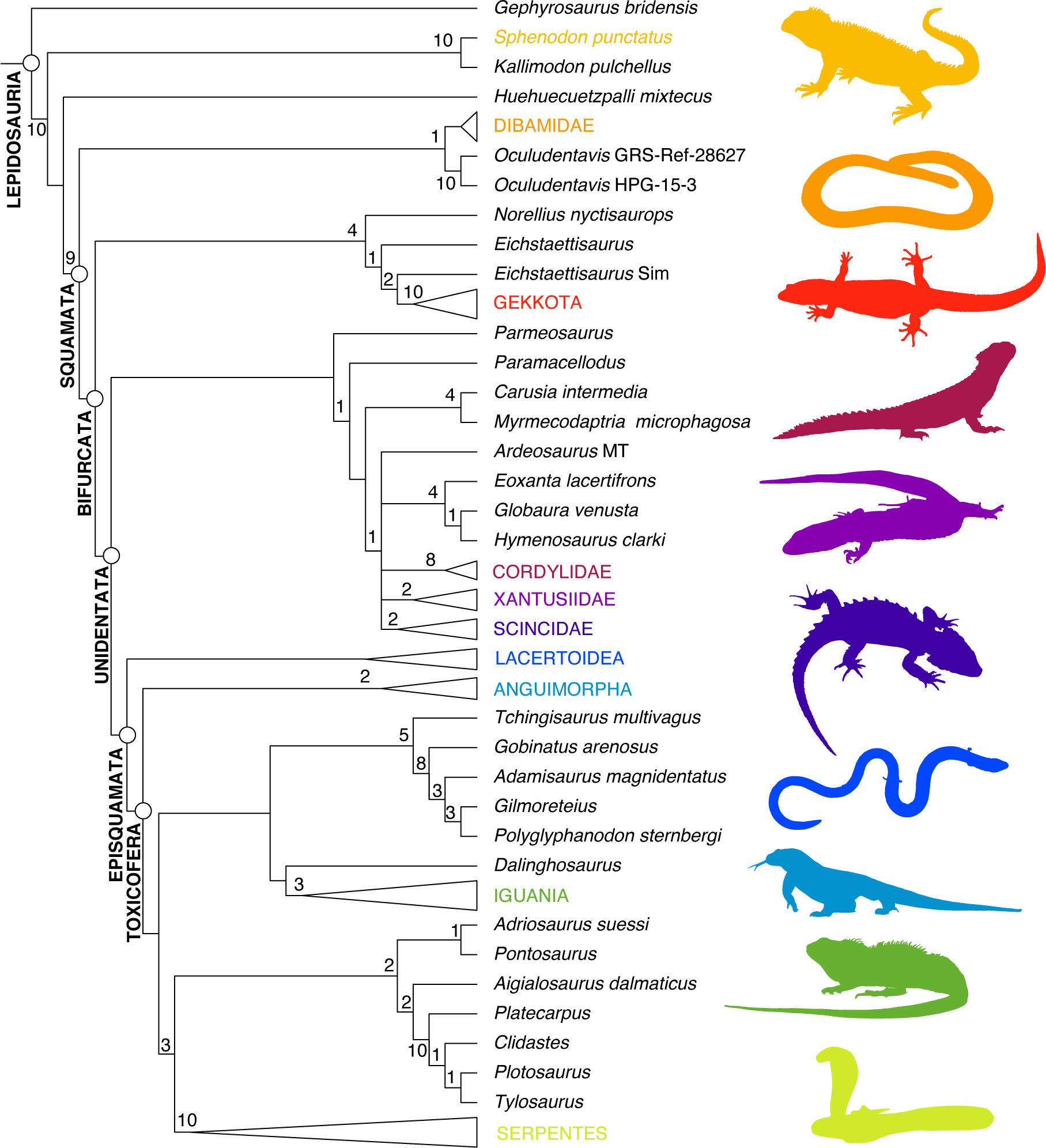
Phylogenetic position of *Oculudentavis* using a lepidosaurian data set (Gauthier et al., 2012) combined with a molecular data set (17), treating some morphological characters as ordered. Crown groups were collapsed, and are represented by silhouettes. *Sphenodon punctatus*, *Anelytropsis papillosus* (Dibamidae), *Sphaerodactylus klauberi* (Gekkota), *Smaug giganteus* (Cordylydae) *Xantusia vigilis* (Xantusiidae), *Tribolonotus gracilis* (Scincidae), *Bachia flavescens* (Lacertoidea), *Varanus komodoensis* (Anguimorpha), *Physignathus cocincinus* (Iguania), *Ophiophagus hannah* (Serpentes). Node values indicate Bremer support.

**Supplementary figure 6.**
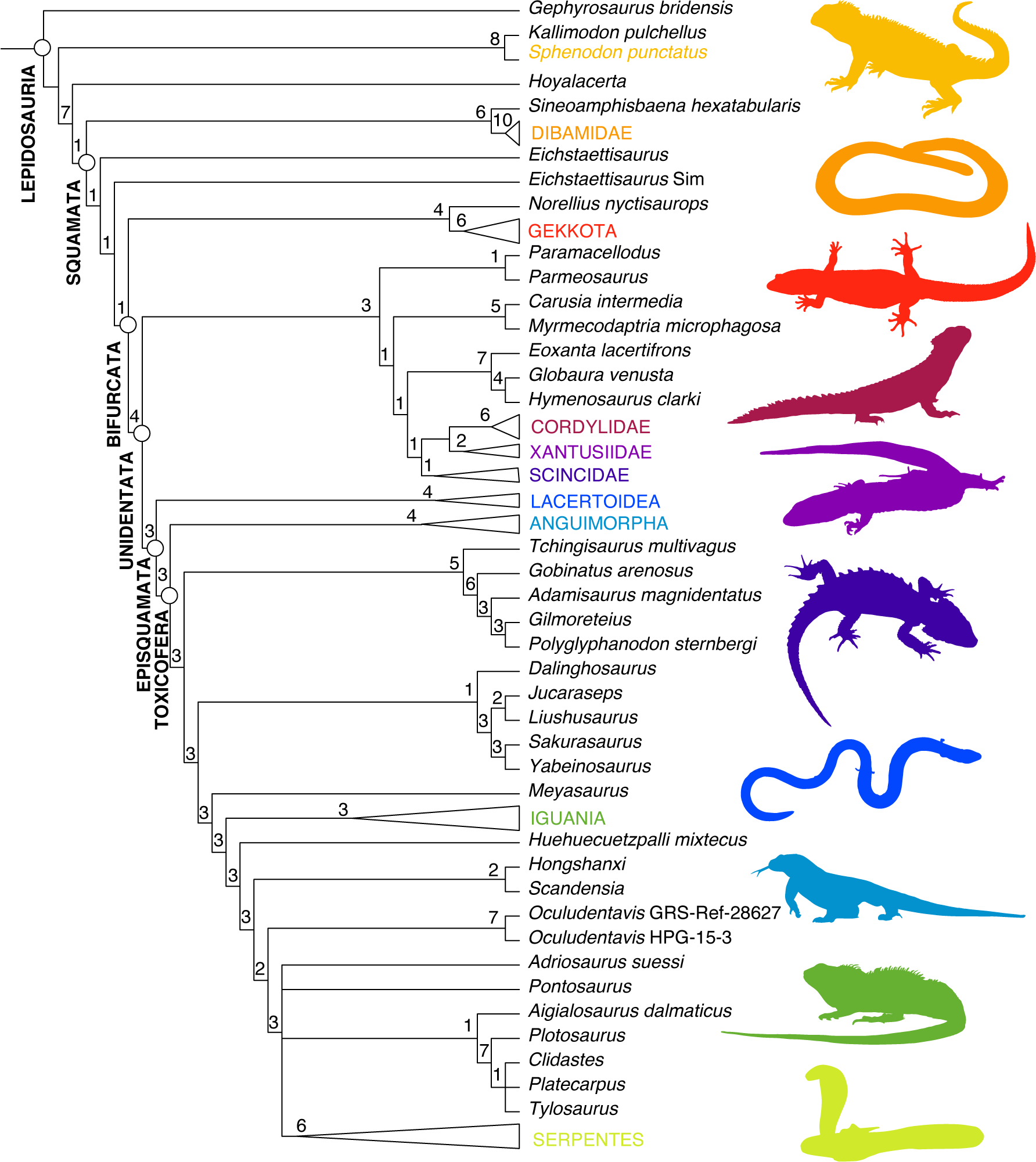
Phylogenetic position of *Oculudentavis* using a lepidosaurian data set (Gauthier et al., 2012) combined with a molecular data set (17), treating all characters as unordered. Crown groups were collapsed, and are represented by silhouettes. *Sphenodon punctatus*, *Anelytropsis papillosus* (Dibamidae), *Sphaerodactylus klauberi* (Gekkota), *Smaug giganteus* (Cordylydae) *Xantusia vigilis* (Xantusiidae), *Tribolonotus gracilis* (Scincidae), *Bachia flavescens* (Lacertoidea), *Varanus komodoensis* (Anguimorpha), *Physignathus cocincinus* (Iguania), *Ophiophagus hannah* (Serpentes). Node values indicate Bremer support.

**Supplementary figure 7.**
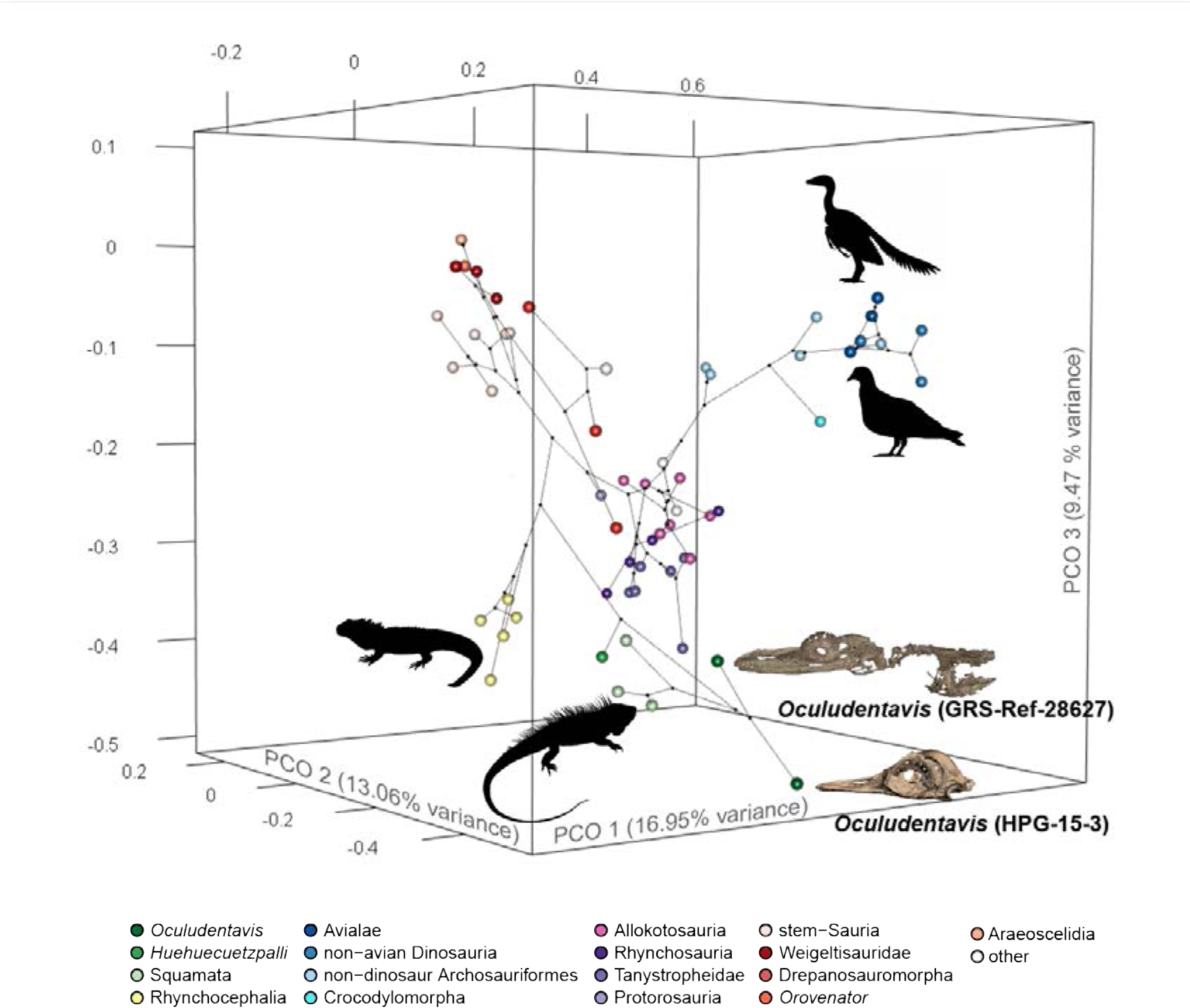
Screenshot of 3D phylomorphospace using the amniote matrix (6).

**Supplementary figure 8.**
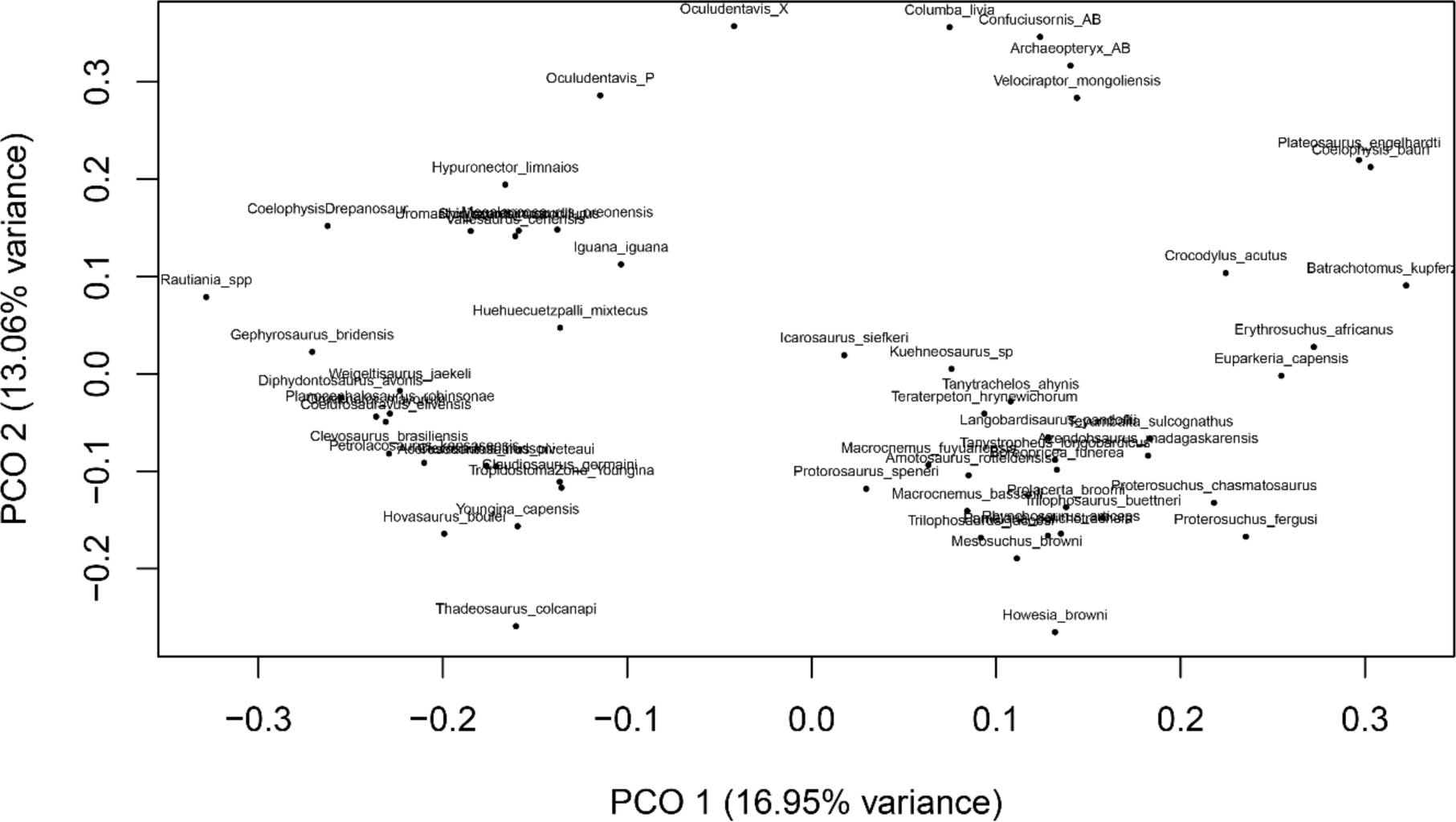
2D morphospace using the amniote matrix (Pritchard and Nesbitt, 2017).

**Supplementary figure 9.**
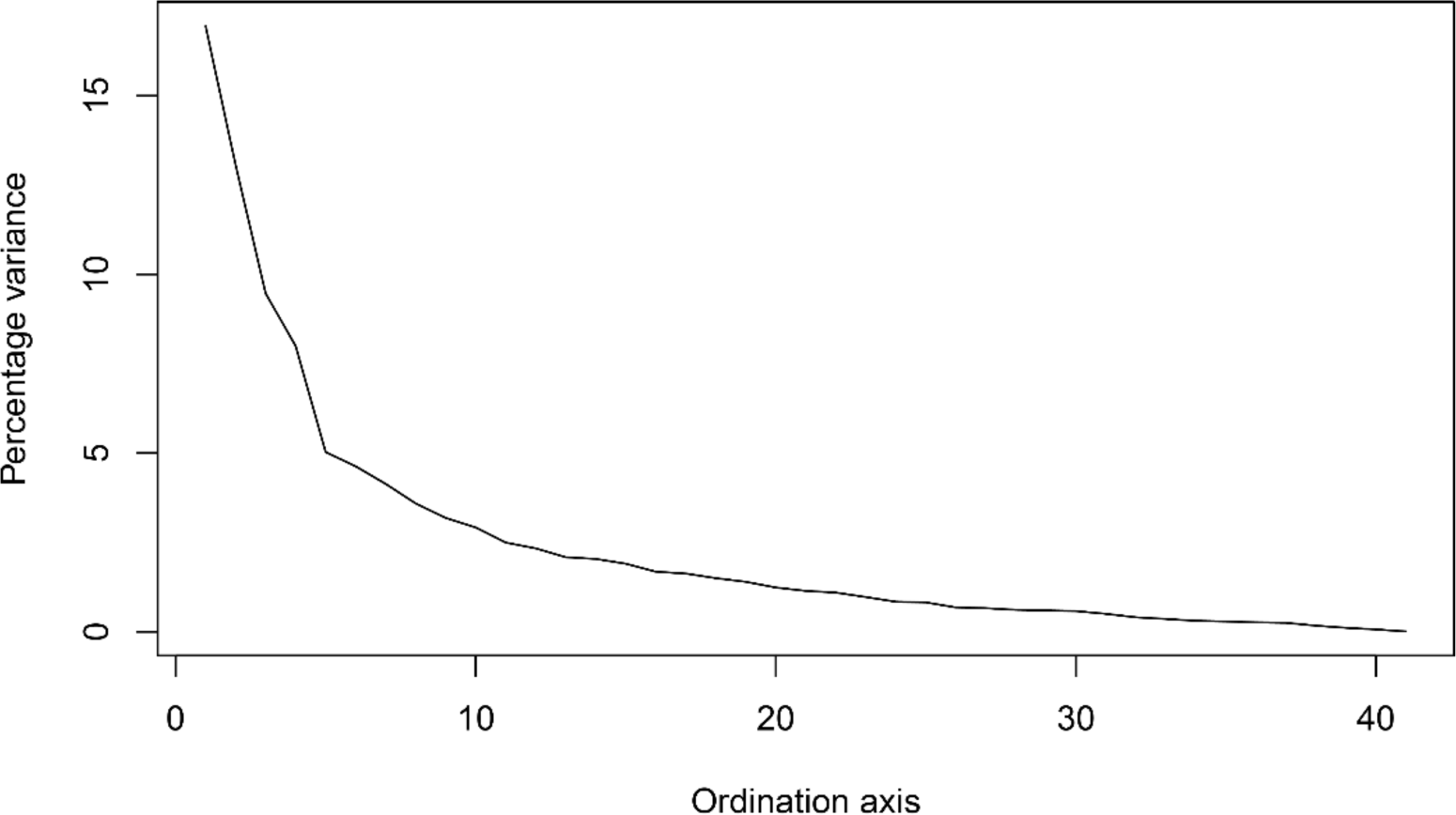
The scree data plot, showing variance explained by each PCO.

**Supplementary figure 10.**
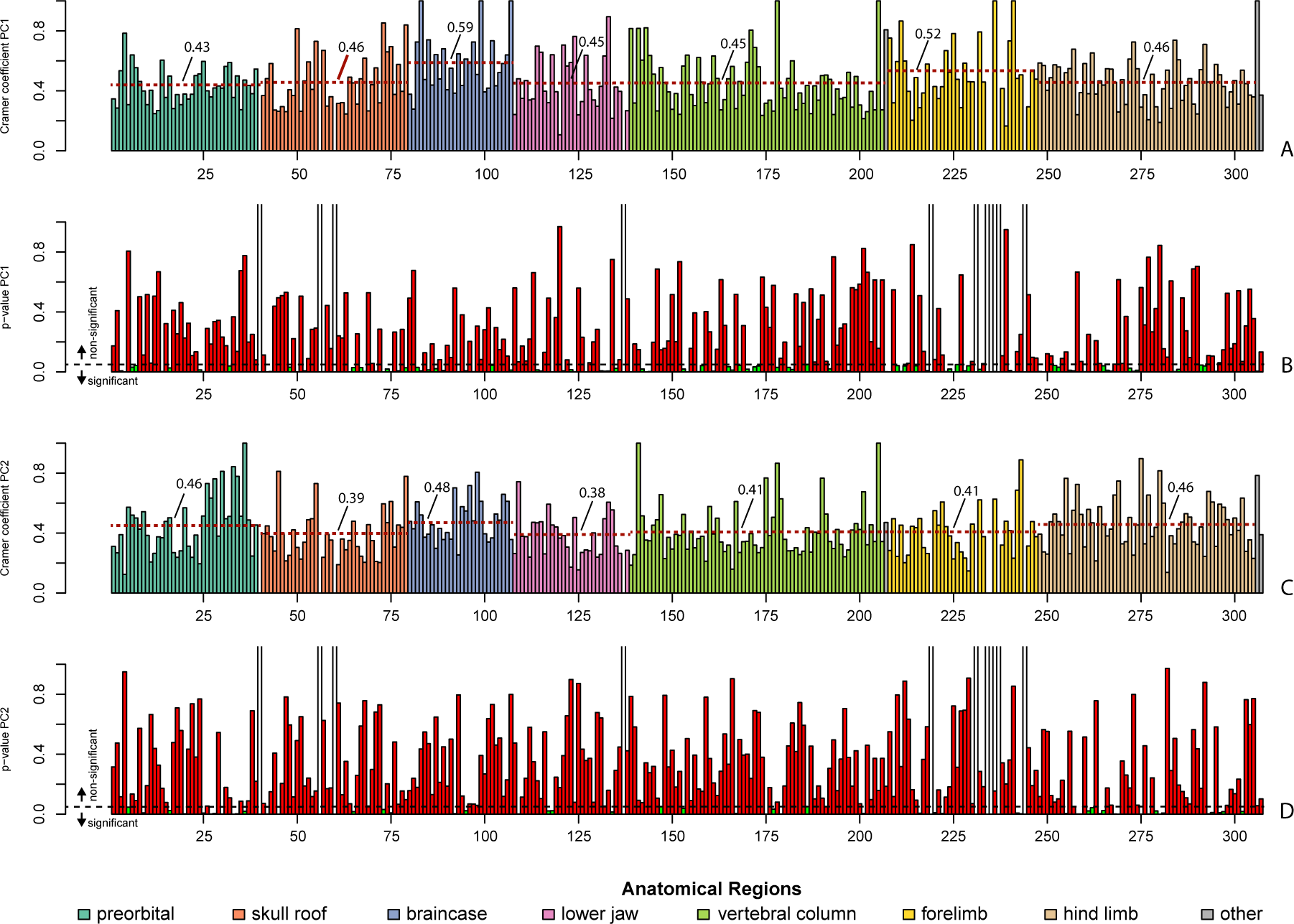
Cramér coefficients and p-values for PC1 and PC2.

**Supplementary figure 11.**
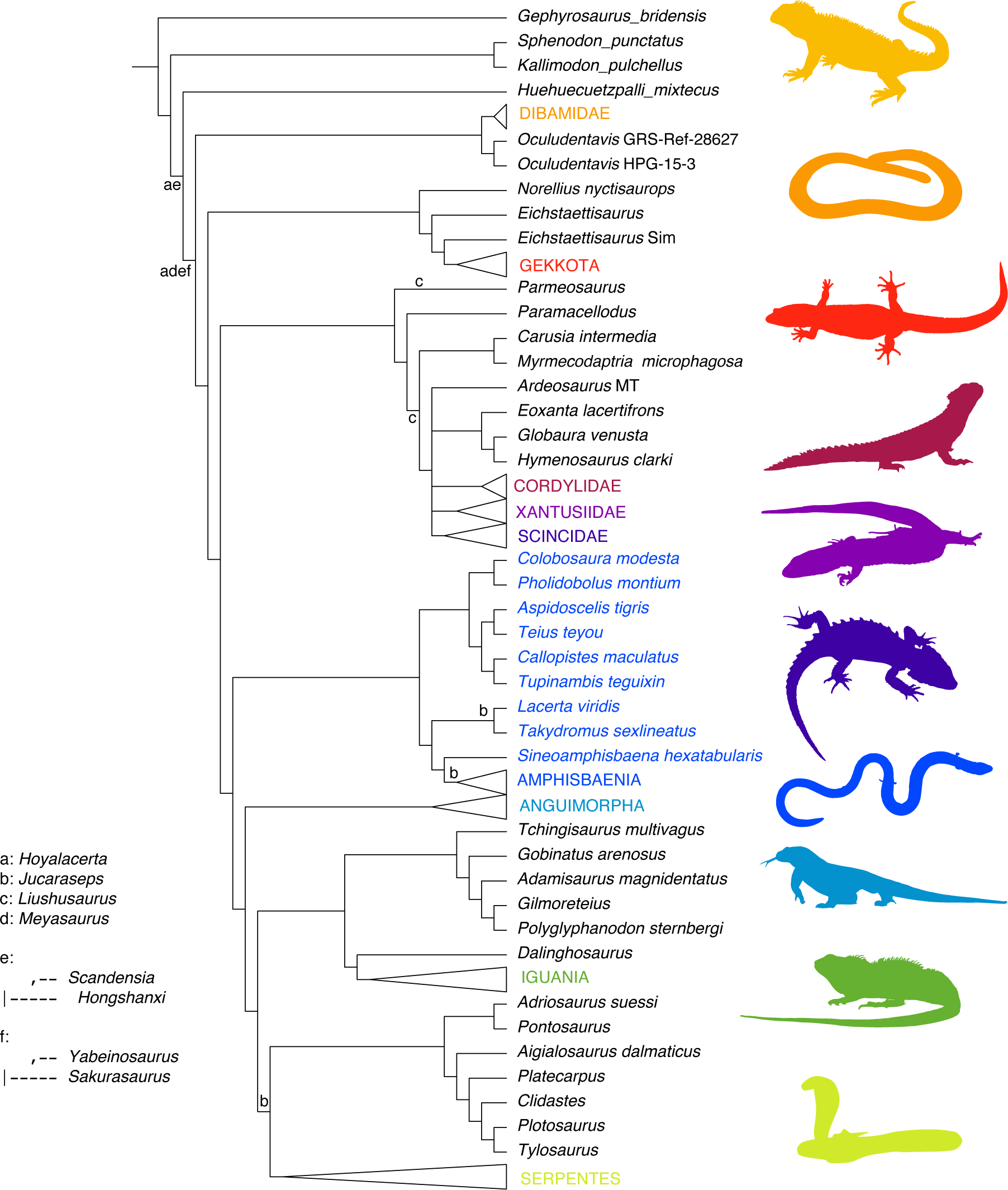
Phylogenetic position of taxa identified as wildcard taxa. Tree was built using a lepidosaurian data set (Gauthier et al., 2012) combined with a molecular data set (17), treating some morphological characters as ordered. Crown groups were collapsed, and are represented by silhouettes. *Sphenodon punctatus*, *Anelytropsis papillosus* (Dibamidae), *Sphaerodactylus klauberi* (Gekkota), *Smaug giganteus* (Cordylydae) *Xantusia vigilis* (Xantusiidae), *Tribolonotus gracilis* (Scincidae), *Bachia flavescens* (Lacertoidea), *Varanus komodoensis* (Anguimorpha), *Physignathus cocincinus* (Iguania), *Ophiophagus hannah* (Serpentes **Source for silhouettes used in** Figure 4C. All silhouettes from Phylopic: Silhouette for Crocodylomorpha corresponds to *Crocodylus porosus* (instead of *Crocodylus acutus*) by Steven Traver. No copyright, This image is available for reuse under the Public Domain Dedication 1.0 license. http://phylopic.org/image/a796420f-5158-4e96-acbe-e30ef36b26d6/ Silhouette for *Archaeopteryx* corresponds to *Archaeopteryx lithographica* by Dann Pigdon. No copyright, This image is available for reuse under the Public Domain Dedication 1.0 license. http://phylopic.org/image/74fce2bf-3fc0-497b-9083-b6f600a697fe/ Silhouette for non-avian dinosaurs corresponds to *Plateosaurus engelhardti* by Andrew Knight. This image is available for reuse under the Creative Commons Attribution 3.0 Unported license. http://phylopic.org/image/f6166a6b-10d1-4c6d-95e6-cf37f5300ae5/ Silhouette for crown-birds corresponds to *Columba livia domestica* by Andreas Plank, based on a photo by Luc Viatour. This image is available for reuse under the Creative Commons Attribution 3.0 Unported license. http://phylopic.org/image/a62a398d-793c-48cf-9803-e52118a28639/ Silhouette for squamates corresponds to *Iguana iguana* by Jack Mayer Wood. This image is available for reuse under the Public Domain Dedication 1.0 license. http://phylopic.org/image/5dec03d9-66a2-4033-b1a9-6dbb3485199f/ Silhouette for rhynchocephalians corresponds to *Sphenodon punctatus* (not sampled), by Benchill. This image is available for reuse under the Creative Commons Attribution 3.0 Unported license. http://phylopic.org/image/bb16d7af-7d0b-43db-b44e-22a1a564bf49/

**Supplementary Table 1.**
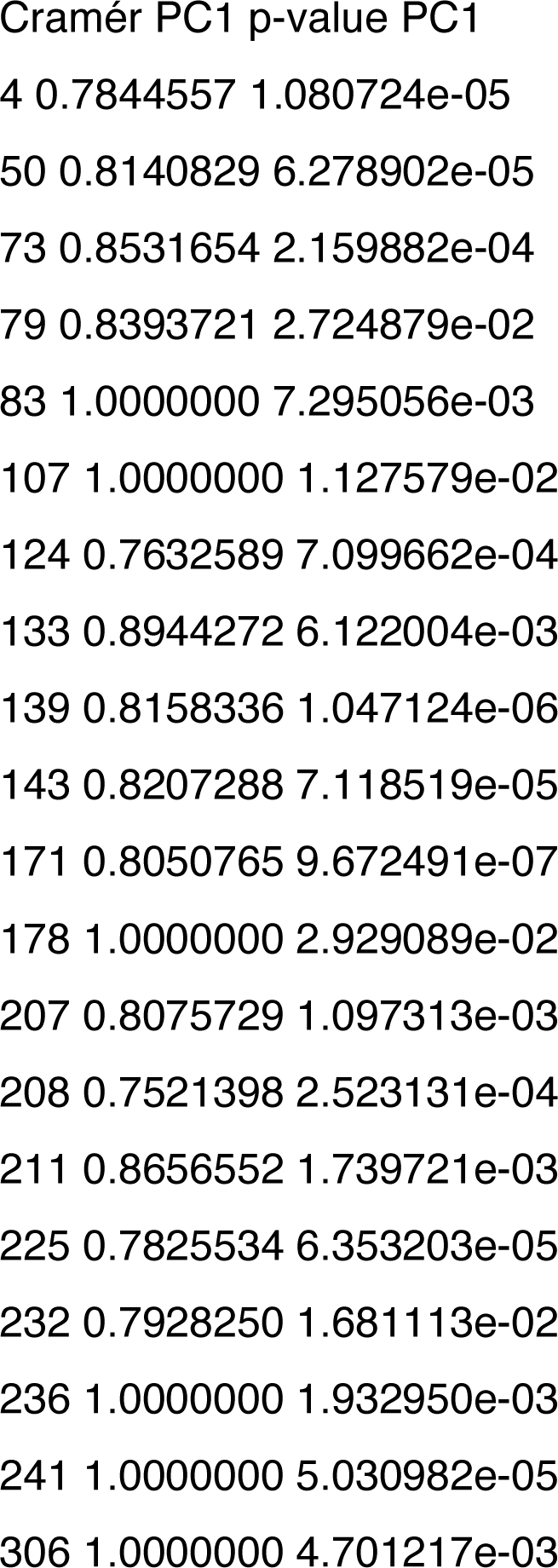
Characters presenting a high Cramér coefficient (<0.75), and p-value < 0.0, for PC1.

**Table S2.** Characters presenting a high Cramér coefficient (<0.75), and p-value < 0.05, for PC2.

|  |  |  |
| --- | --- | --- |
| 28 | 0.7617141 | 4.527297e-03 |
| 30 | 0.8115511 | 3.059130e-04 |
| 33 | 0.8431396 | 7.384333e-05 |
| 34 | 0.7786206 | 1.954688e-03 |
| 36 | 1.0000000 | 1.831564e-02 |
| 45 | 0.8122329 | 1.640635e-06 |
| 175 | 0.7677472 | 1.669355e-04 |
| 243 | 0.8888194 | 2.321543e-03 |
| 255 | 0.7657438 | 4.218288e-03 |
| 275 | 0.8975275 | 1.139072e-05 |
| 280 | 0.8153439 | 3.274084e-05 |

**Table S3.**
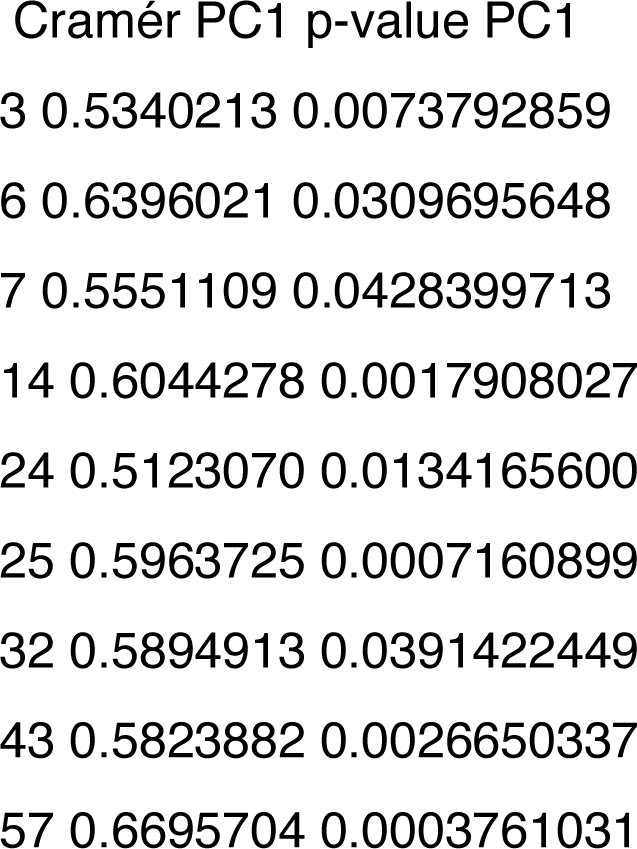
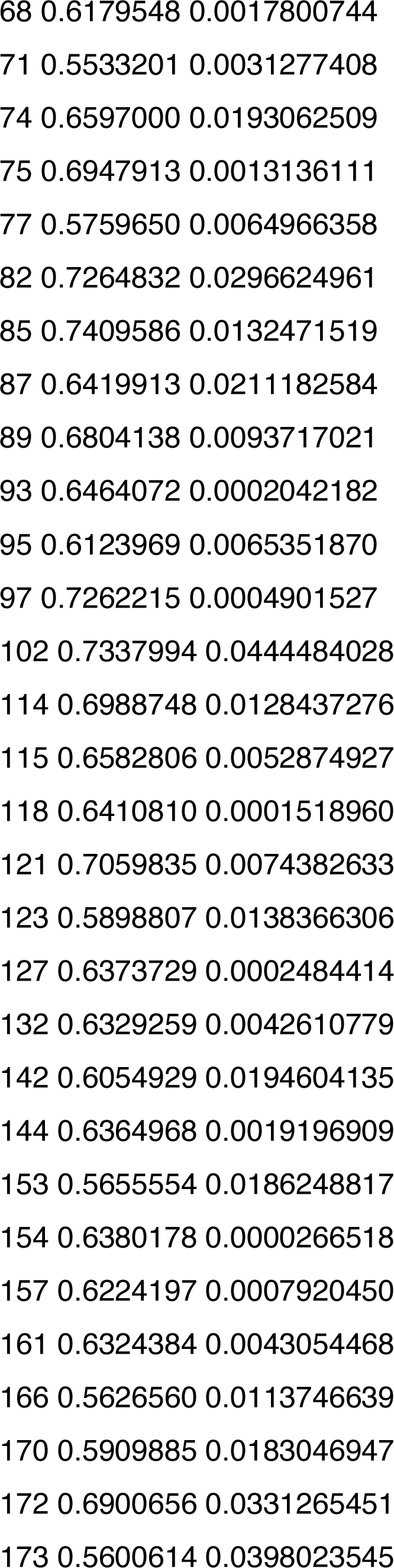
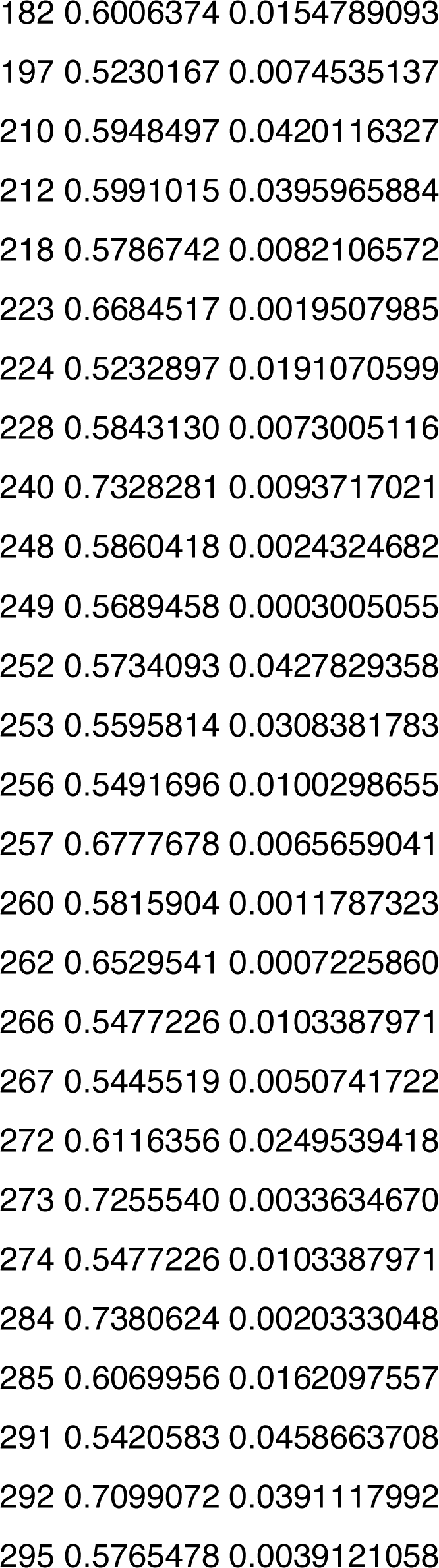
Characters presenting a moderate Cramér coefficient (0.5<x<0.75), and p-value < 0.0, for PC1.

**Table S4.**
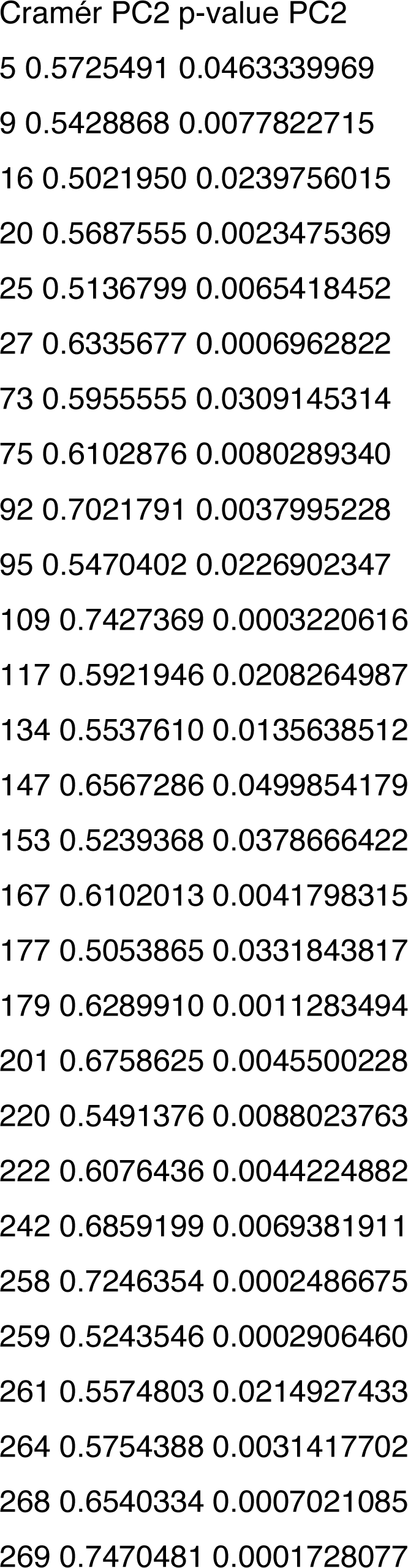
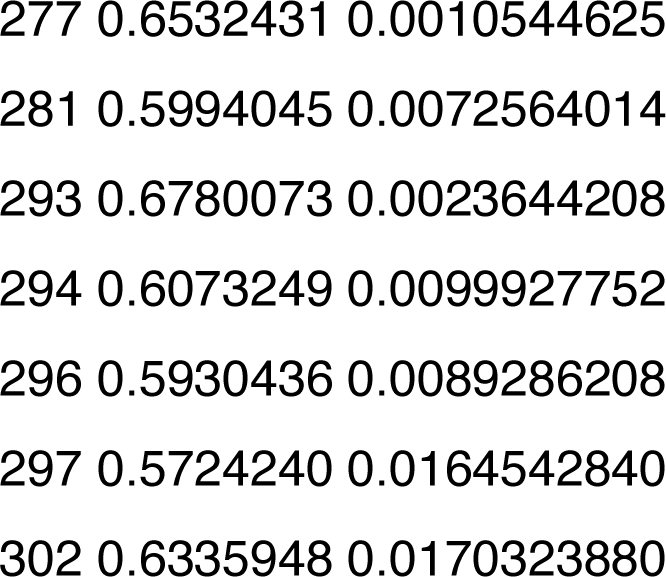
Characters presenting a moderate Cramér coefficient (0.5<x<0.75), and p-Cramér PC2 p-value PC2

